# Insights into Temporal and Spatial Dynamics of Short Association Fiber Formation in the Human Fetal Brain

**DOI:** 10.1101/2025.10.20.683525

**Authors:** Bo Li, Željka Krsnik, Lana Pierotich, Ivica Kostović, Simon K. Warfield, P. Ellen Grant, Davood Karimi

**Affiliations:** Computational Radiology Laboratory, Boston Children’s Hospital, Boston, MA, USA; Harvard Medical School, Boston, MA, USA; Croatian Institute for Brain Research, School of Medicine, University of Zagreb School of Medicine, Zagreb, Croatia; Fetal Neonatal Neuroimaging and Developmental Science Center, Boston Children’s Hospital, Boston, MA, USA; Division of Newborn Medicine, Boston Children’s Hospital, Boston, MA, USA

## Abstract

Short association fibers (SAFs) form the local scaffold of cortical connectivity, supporting early functional specialization and marking sites of neurodevelopmental vulnerability. However, their development before birth remains largely unknown. Leveraging advanced fetal diffusion MRI and a histology-validated framework, we present the first in-utero reconstruction of SAF pathways in the human brain. We tracked their volumetric and microstructural developmental trajectories in 243 fetuses spanning a critical period when the brain’s connectome is rapidly forming. We found that SAFs emerge before sulcal folding, initially as flat, loosely arranged pathways along the subplate–white matter interface, and later reorganize into coherent U-shaped bundles. Their maturation followed a sensorimotor-to-association gradient, paralleling cortical development. Nonlinear, bundle-specific trajectories captured multiphasic maturation, with subplate dynamics preceding increases in axonal coherence and early myelination. By filling a missing link in lifespan brain connectivity, this study establishes a prenatal reference for cortical wiring and provides tools for investigating the origins of neurodevelopmental disorders.

## 1 Introduction

Short association fibers (SAFs) are a dense network of curved axonal pathways that interconnect neighboring gyri within the same cerebral hemisphere, forming the fine-scale infrastructure of local cortical communication. These fibers, historically described as Arnold’s or Meynert’s arcuate fibers, or as U-fibers due to their arched trajectories around sulcal walls (Arnold, 1838; Meynert et al., 1872; Schmahmann and Pandya, 2007), contribute to the integrity of cortical circuits by mediating short-range communications between sets of neurons in different functionally specified cortical gyri. Unlike long-range projection, commissural, and association fibers that coordinate distant brain regions, SAFs support regionally confined integration and have been implicated in processes such as cortical specialization, plasticity, and localized developmental tuning (Yoshino et al., 2020; Movahedian Attar et al., 2020). Though they constitute a substantial portion of the white matter (WM) volume and likely underpin key aspects of early circuit formation, SAFs remain understudied, in part due to their small size, high individual variability (Román et al., 2022), and limited visibility in low-resolution diffusion MRI reconstructions. These challenges are magnified in the fetal brain, where transient developmental zones, ongoing cellular migration, and rapid changes in both tissue composition and cytoarchitecture complicate anatomical interpretation. Fetal diffusion MRI is further hindered by motion artifacts and limited diffusion encoding schemes. Together, these factors obscure the observation of emerging SAF systems during this critical period of development.

Human fetal brain development is characterized by rapid, regionally differentiated changes in tissue composition (histogenesis), progressive establishment of corticocortical connectivity, and a developing cortex-WM interface shaped by structural and molecular changes such as subplate dynamics, laminar differentiation, and gyrification (Kostović et al., 2014a). Of particular interest is the distal (superficial) compartment of the developing cerebral wall, comprising the cortical plate (CP), gyral WM, and the subplate, a transient zone interposed between them (Kostović et al., 2014b). The subplate is the most prominent lamina of the fetal brain and persists into the early postnatal period. It serves as a site of the earliest synaptic cortical activity and a growth zone for emerging corticocortical connections, whose large size in humans has been linked to the increased number and variability of corticocortical connections compared to other mammals (Kostovic and Rakic, 1990; Bystron et al., 2008). To describe the developing fetal WM, we adopted a classical five-segment division (von Monakow, 1905): (I) periventricular zone, (II) sagittal strata and crossroads, (III) centrum semiovale, (IV) gyral WM, and (V) intracortical fiber zone. As development progresses, the abundant extracellular matrix (ECM) in the subplate diminishes, its deeper territory is progressively occupied by the expanding centrum semiovale and future gyral WM (segments III-IV), while cortical afferent fibers that had temporarily waited in the subplate relocate into the overlying CP (Kostović et al., 2014a).

Additionally, the fetal brain undergoes a progressive shift in fiber-architectonic arrangement during the second half of gestation (20-40 weeks), from predominantly radial and tangential organizations, to increasingly multidirectional organizations characteristic of emerging corticocortical connectivity (Takahashi et al., 2012; Xu et al., 2014; Huang and Vasung, 2014). During this time, projection pathways are largely in place, while long-range commissural and associative pathways, along with SAFs, develop in a temporally sequential but substantially overlapping manner, though all fiber systems continue to mature postnatally (Burkhalter et al., 1993; Hevner and Kinney, 1996). This sequence, from deep long-range systems to progressively refined superficial local connections, reflects the hierarchical assembly of structural connectivity in the developing fetal brain.

However, knowledge of the developing SAFs, major components of the local circuits, remains limited, as they are difficult to trace and visualize in the human fetal brain. Even experimental animal models have been largely unsuccessful in demonstrating the trajectories of developing fiber pathways, with the notable exception of studies in fetal monkeys that used invasive prenatal access and autoradiographic tracers to reveal select cortical connections (Rakic, 1976; Goldman-Rakic, 1987). In vivo visualization using tractography offers a noninvasive alternative for investigating this important fiber system. However, existing fetal brain studies have focused exclusively on long-range pathways and reconstructed only a small number of major tracts (Kasprian et al., 2008; Jaimes et al., 2020; Wilson et al., 2021; Calixto et al., 2025b). SAFs, by contrast, are short, spatially heterogeneous, and embedded within the fluid-rich subplate zone near the rapidly folding CP. Their small size, coupled with the small physical dimensions of the fetal brain itself, makes these fibers especially difficult to resolve using existing tractography methods. Moreover, most existing WM atlases and fiber bundle parcellation strategies are designed for large, major bundles and depend on adult-derived regions of interest (ROIs) and fixed-depth sampling approaches (Oishi et al., 2009; Catani et al., 2012; Guevara et al., 2012; Rojkova et al., 2016), which do not generalize to the rapidly changing organization of the developing fetal brain. As a result, fundamental questions remain unresolved: When do SAFs first exhibit a coherent structure? How do they develop in size, orientation, and microstructural organization throughout gestation?

In light of these gaps, we developed a histology-informed, gestational-age-adaptive pipeline for reconstructing and quantifying SAFs during the second and third trimesters (Figure 7). Our approach incorporates age-adaptive tissue priors, optimized tractography for SAFs and fetal diffusion MRI, and validation with gestational age-matched postmortem histological data to ensure anatomical fidelity. We applied this pipeline to a multi-shell dataset of 243 normally developing fetuses between 22 and 38 gestational weeks (GW), enabling robust identification of over 250 reproducible SAF bundles confined to the superficial compartment, while excluding intermediate and deep zones such as WM segments I-III (Figure 8). We characterized the spatiotemporal development of SAF pathways using volume and free-water-eliminated diffusion tensor imaging (DTI) metrics. This study provides the first whole-brain, gyral-level characterization of SAF development in utero, establishing a spatiotemporal reference for the maturation of short-range connectivity during the second and third trimesters. These findings not only fill a longstanding gap in our understanding of the fetal connectome but also lay the foundation for normative baselines essential for identifying atypical neurodevelopment. By making the framework and quality-controlled results publicly available, this work offers a scalable framework to map human brain connectivity before birth, advancing both basic neuroscience and clinically translational research beyond existing postnatal preterm models and opening new avenues to investigate the origins of neurodevelopmental vulnerability in vivo.

## 2 Results

### Successful reconstruction of SAFs in utero

To systematically chart the spatiotemporal development of SAFs across the fetal cortex, we analyzed high-resolution, multi-shell fetal diffusion MRI data (N=243; age: 22-38 GW), provided by the Developing Human Connectome Project (dHCP^1^). This research dataset offers the highest available spatial and angular resolution for in-utero diffusion imaging, enabling the reconstruction of SAFs that remain inaccessible with clinical-quality data. We developed a fully automatic, fetal-optimized analytical pipeline tailored to the challenges of SAF reconstruction, informed by developmental neuroanatomy. Whole-brain fiber pathways were reconstructed using anatomically constrained probabilistic tractography (Tournier et al., 2010), seeded from a curated CP-subplate interface, which enabled pathway continuity through regions where CP over-segmentation would otherwise obstruct propagation, such as at the gyral crown or beneath the insular cortex. To accommodate the diverse geometries of developing fibers and the rapidly changing microstructure across gestation, we implemented an ensemble tracking strategy with ten tractography configurations varying in angular threshold, fiber orientation distribution (FOD) amplitude cutoff, and curvature sensitivity. The resulting ensemble included one million streamlines per subject. To isolate SAFs, we applied an age-adaptive superficial compartment mask that included the subplate and emerging gyral WM but excluded streamlines traversing or terminating in deeper compartments, including the intermediate zone (long-range association fibers), periventricular zone (commissural fibers), and subcortical or brainstem regions (projection fibers). Anatomical definition of this mask was guided by the laminar organization of the cerebral wall (Kostović et al., 2014b; von Monakow, 1905; Judaš et al., 2013) and the CRL fetal brain atlas (Gholipour et al., 2017). For fetuses under 32 weeks, the subplate was delineated directly from the atlas. For later gestation, when the subplate becomes indistinct, we optimized the mask through varying degrees of WM erosion followed by an iterative, inside-out “peeling” procedure to conservatively restrict streamlines to the superficial compartment. For example, beneath the insular cortex, SAFs were constrained to remain above the external capsule, the outermost deep WM structure (Kostović et al., 2014a).

The resulting tractograms achieved high cortical coverage and, for the first time, enabled in-utero visualization of emerging SAFs during early stages of development. Unlike adult SAFs, which are often referred to as U-fibers, fetal SAFs did not initially exhibit this arcuate configuration (Figure 1a). Instead, early pathways appeared as flat structures that gradually reshaped into curved trajectories, paralleling sulcal deepening over time (see also the bundle-wise visualization, e.g., Figure 5). This morphological evolution followed a sensorimotor (central) to peripheral spatial progression, which largely mirrors the known spatiotemporal gradient of cortical gyrification (Chi et al., 1977; Dubois et al., 2008; Yun et al., 2022). For instance, bundles surrounding the central sulcus exhibited early curvature, followed by those in the inferior frontal (including Broca’s area) and superior temporal gyri (including Heschl’s gyrus), and later by bundles in higher-order associative regions such as the inferior temporal and superior/middle frontal gyri. By 35 GW, widespread U-fibers and coherently oriented SAF bundles were observed (Figure 1c), similar to prior observations at term age (42 GW) (Takahashi et al., 2012).

**Figure 1.**
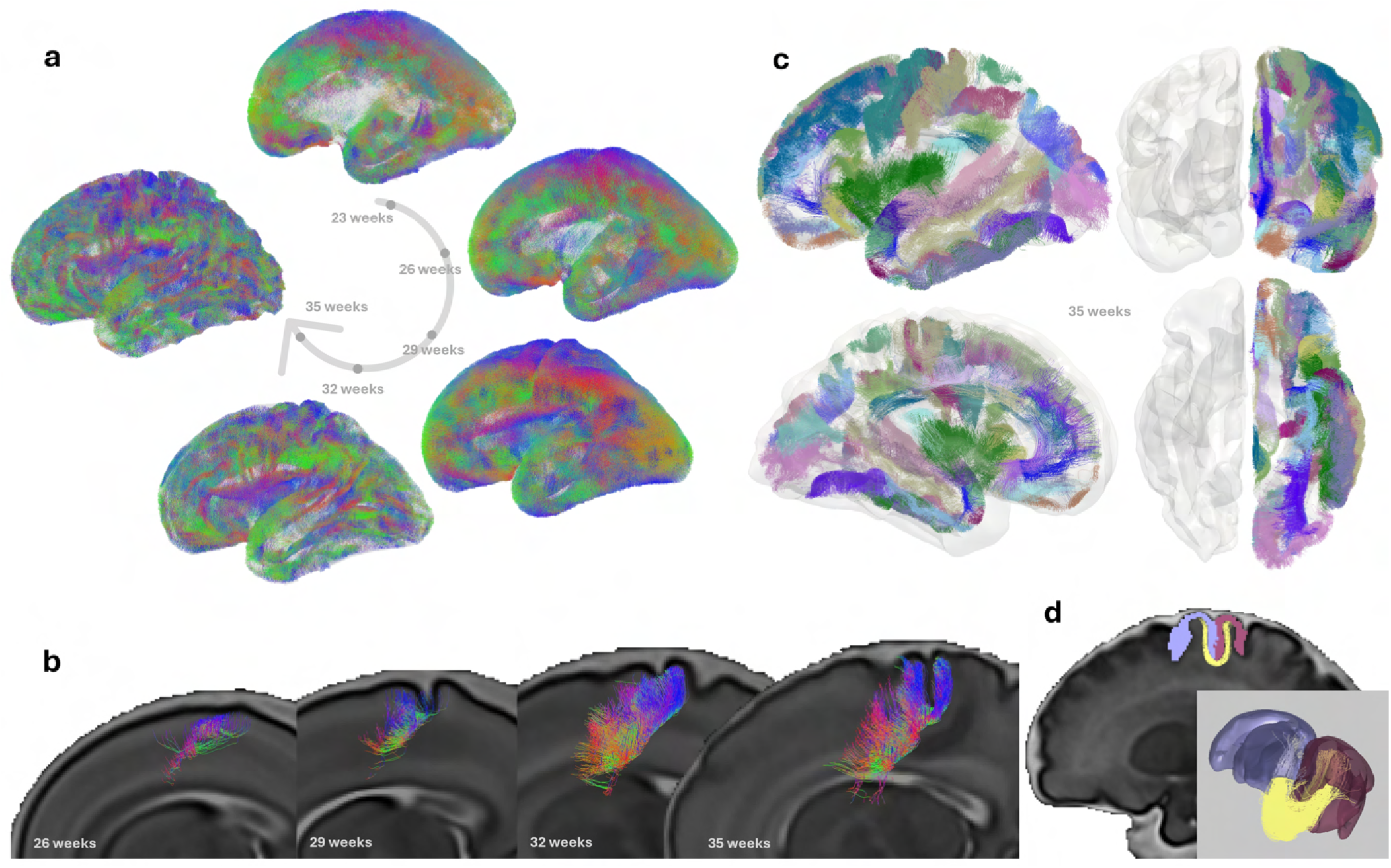
Progression of short association fibers (SAFs) formation across the human cortex, revealed by in-utero diffusion MRI. **a**. Whole-brain tractograms at five key gestational stages (23, 26, 29, 32, and 35 weeks) demonstrate the spatiotemporal development gradient of SAFs in the fetal brain. The morphological changes of the SAF system were closely related to cortical gyrification, which begins from the central sulcus outward to the poles. These reconstructions, generated using a fetal-specific processing pipeline and data from the dHCP diffusion MRI, capture for the first time the emergence of SAF systems in utero. **b**. Focusing on the precentral-postcentral connection, we illustrate a developmental transformation of arcuate SAF geometry over time. To visualize this evolution, 3D fiber bundles are overlaid on 2D sagittal T2w MRI slices (left view), which provides anatomical context, including changes in gyrification and cerebral wall development. The background slice represents a medial plane and does not capture the lateral cortical surface; thus, some SAFs may appear to traverse the ventricular region due to the projection of 3D bundles onto the 2D image. At 26 GW, this inter-gyri connection appears as a loosely organized structure with 2-3 dominant orientations, including radial (blue-purple) and tangential (green) trajectories. The presence of multiple orientations at earlier stages may reflect the plexiform organization of various contingents of growing axons within the developing subplate (Kostovic and Rakic, 1990). As development progresses, fiber pathways present an anterior-posterior alignment, running parallel to one another along the medial-lateral axis of the sulcal base. By 32 GW, the tract adopts a curved trajectory at both ends, wrapping around the deepening sulcal walls. This configuration sharpens over time, forming a coherently organized and sharply delineated U-shaped bundle by 35 GW. **c**. By 35 GW, widespread SAFs with canonical U-shaped geometry are visible throughout the cortex, resembling adult-like short-range connectivity. Shown are 3D renderings from lateral, anterior, inferior, and medial (left hemisphere) views. **d**. Streamlines were grouped into candidate bundles based on their endpoints, each linking a unique pair of adjacent gyri or distinct regions within the same gyrus. The illustrative results in this figure were performed on the dHCP fetal template (Uus et al., 2023) using the proposed fetal-specific analysis pipeline. Fiber orientation in (**a, b**) follows standard diffusion tensor imaging color coding: red = mediolateral, green = anteroposterior, and blue = superoinferior.

A particularly dynamic transformation in the geometry of SAF bundles was observed, illustrated here by the precentral-postcentral connection (Figure 1b). At 26 GW, this inter-gyri bundle appears as a loosely-organized structure with 2-3 dominant orientations, involving radial (likely reflecting neuronal migration pathways (Rakic, 1972)) and tangential trajectories. Between 29 and 32 GW, these early, partially intersecting pathways progressively reorganized into more coherent, compact, and arcuate forms that delineate the deepening sulci along the subplate-WM interface. The presence of multiple orientations and diffuse geometry at early stages is possibly related to the underlying complexity of the subplate zone, characterized by a water-rich ECM, various migrating and developing cells, and a plexiform meshwork of growing axons with variably oriented neuronal processes (Kostovic and Rakic, 1990). The observed orientation clusters may represent a macroscopic signature of this temporally dynamic and structurally complex axonal environment within the developing subplate. This flat-to-arcuate transition between 29 and 32 weeks occurred within the broader framework of strata-like ingrowth of corticocortical fibers, reflecting a progression toward spatially confined and directionally coherent bundles. By 35 GW, these pathways increasingly adopted the characteristic U-shape commonly observed in the adult brain.

### Histological validation of fetal SAF reconstructions

To establish the biological fidelity of the in utero reconstructed SAFs, we cross-validated them against postmortem histological sections from human fetal brains. These comparisons also informed interpretation of SAF architecture within its laminar context, using histological markers. Acetylcholinesterase (AChE) histochemistry reveals key features of the developing laminar organization of the fetal cerebral wall, with intense reactivity highlighting the subplate and CP. These histological landmarks informed the design of the superficial compartment mask, defined on T2-weighted MRI, to isolate SAFs from whole-brain pathways. MRI–AChE staining correlations along the developing cerebral wall are illustrated at 28 and 36 GW (Figure 8). For direct comparison on fiber architecture, we focused on later gestational stages (35-40 GW), where myelin- and subplate-associated markers are available through Myelin Basic Protein (MBP) and Chondroitin Sulfate (CS-56) staining. In-utero diffusion MRI results at 35 GW were selected to match the anatomical levels of the MBP and CS-56 staining, ensuring consistency in regional anatomy and developmental stage.

In the central region at 35 GW, diffusion MRI consistently revealed SAF pathways bridging the precentral and postcentral gyri (Figure 2**a-b**). These reconstructed streamlines showed precise spatial correspondence with MBP-stained SAFs located immediately beneath cortical layer VI. On MBP-stained sections, these SAFs were anatomically distinct from deeper association and projection pathways, confirming their confinement to the superficial compartment. Importantly, although these fibers had not yet developed mature myelin, the associative parietal somatosensory cortex exhibited early myelination of SAFs by 35 GW, supporting their advanced maturation relative to other cortical regions (Kinney et al., 1988).

**Figure 2.**
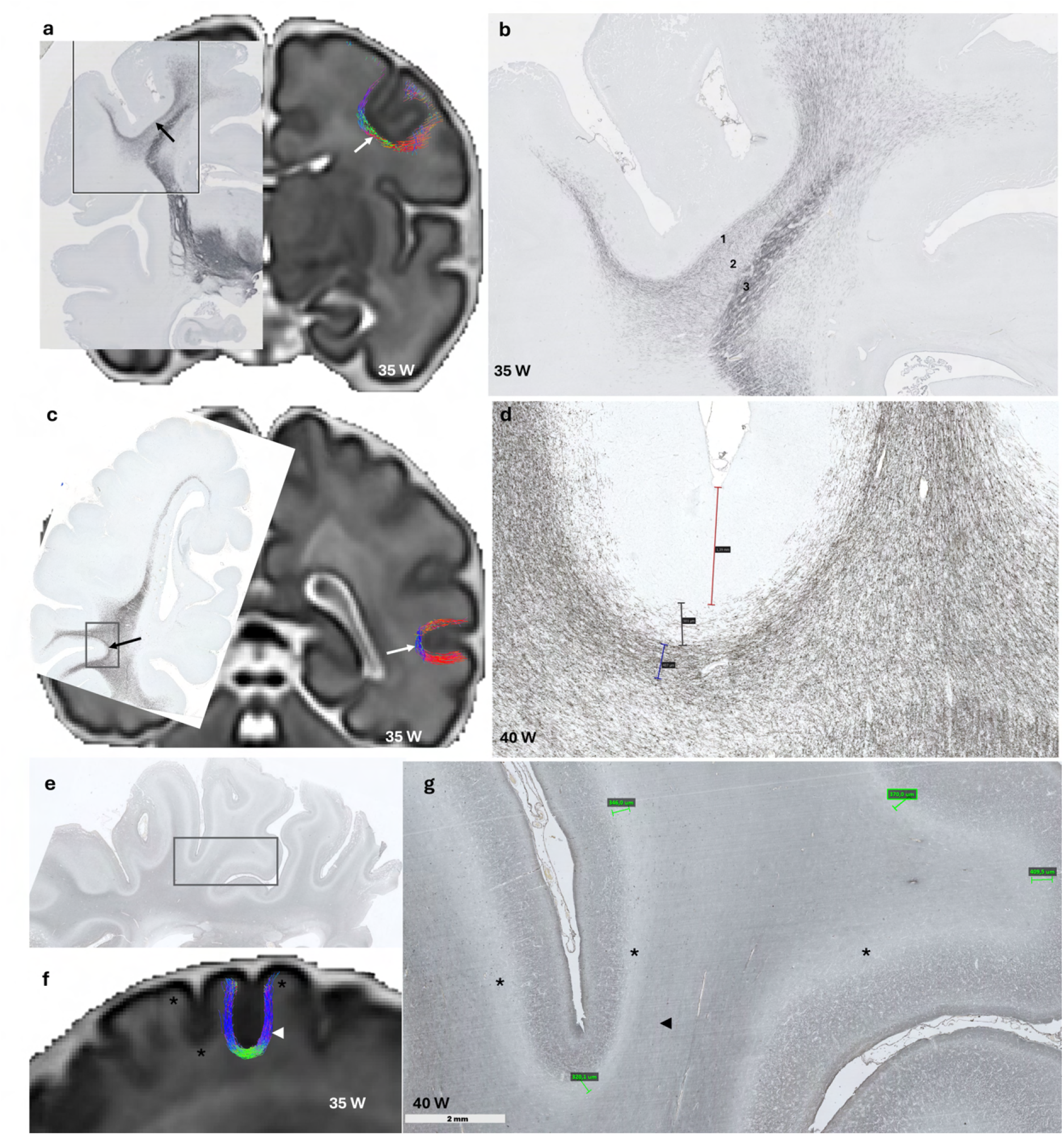
Histological validation of reconstructed short association fiber (SAF) systems through side-by-side comparisons of in-utero MRI findings (**a**,**c**,**f** ) and postmortem histological sections (**b**,**d**,**e**,**g**) at matched anatomical locations, illustrating the axonal organization of SAFs and adjacent cerebral compartments. **a-b**. In the central cortex at 35 GW, in-utero MRI framework identifies U-shaped SAFs (**a**, white arrow) situated directly beneath cortical layer VI and along the wall of the central sulcus, corresponding to myelinated SAFs (**a**, black arrow) marked by Myelin Basic Protein (MBP)-staining on postmortem sections. These SAFs (**b**-1) are distinct from the extension of the immature centrum semiovale which contains long associative pathways (**b**-2), and from the major projection pathways streaming from the internal capsule (**b**-3), highlighting the transient multilaminar organization of fetal cortical connectivity, which evolves dynamically throughout gestation. **c-d**. In the temporal region of a 40 GW brain, MBP staining reveals a tightly bundled SAF system (**c**, black arrow; **d**, blue bracket; *∼* 407 *µ*m), located primarily within gyral white matter while beneath the subplate remnant (SPrm; **d**, gray bracket; *∼* 501 *µ*m) and cortical layers I-VI (**d**, red bracket; *∼* 1.39 mm). The SPrm contains loosely arranged plexiform fibers, but is clearly delineated from the dense U-shaped pathways at sulcal fundi. Oligodendrocytes appear as black dots in **d**. The corresponding SAFs were identified from in-utero MRI at 35 GW (**c**, white arrow). **e-g**. Chondroitin sulfate (CS-56)-stained sections further delineate the SPrm, visible as a pale band in the frontoparietal region (**g**, asterisks). CS-56 labels chondroitin sulfate proteoglycans enriched in the extracellular matrix of the subplate, highlighting its residual presence near term. The SPrm is narrowest at the sulcal fundi (**g**, green bracket; *∼* 320 *µ*m), and becomes progressively thicker in the gyral walls (*∼* 346 *−* 370 *µ*m) and crowns (*∼* 410 *µ*m). During late gestation, the gyral white matter develops, and parallels with the reduction of the subplate. It gradually occupies the former subplate position in the proximal parts of cortical gyri. The primary location of SAFs shifts from within the subplate (**f**, white arrowhead) to the underlying, newly developed gyral white matter (**g**, black arrowhead). Together, these histological observations support the anatomical fidelity of the proposed in-utero MRI-based analytical framework and provide microstructural context for interpreting SAF development, including its spatial relationship with related laminar organizations and fiber systems.

In the temporal lobe at 40 GW, MBP staining identified a clearly delineated SAF layer located predominantly in the newly developing gyral WM, beneath the subplate remanent (SPrm) and cortical layers I-VI (Figure 2**c-d**). In this specimen, the SAF layer measured ∼407 *µ*m in thickness, the SPrm ∼501 *µ*m, and the cortical layers ∼1.39 mm, highlighting the need for sufficient spatial resolution and biological specificity. Additional histological views (Figure 2**g**) revealed ongoing subplate dissolution, with SPrm thickness varying across cortical regions, from ∼320 *µ*m in sulcal depths to *>* 400 *µ*m in gyral crowns. These laminar profiles reflect the ongoing compartmentalization of the fetal cortex and the spatiotemporal development of SAF systems within this context, with changes in the subplate serving as an indirect indicator (Kostović et al., 2014a).

In the frontal lobe, a coronal section at 40 GW revealed multiple SAF systems spanning diverse cortical territories (Figure 3). Myelin-stained sections and in-utero MRI jointly delineated several SAF connections between the middle frontal and the opercular portions of the inferior frontal gyri (Figure 3b), between the superior and middle frontal gyri (Figure 3e), and between the supplementary motor area and the mid-cingulate cortex (Figure 3h). These pathways traversed gyri of varying morphological complexity, including both middle and late folding gyri (Appendix Table A1), and extended across both lateral and medial surfaces. It is additionally important to see from the myelin stains that the frontal associative regions mature not much behind the somatosensory cortex.

**Figure 3.**
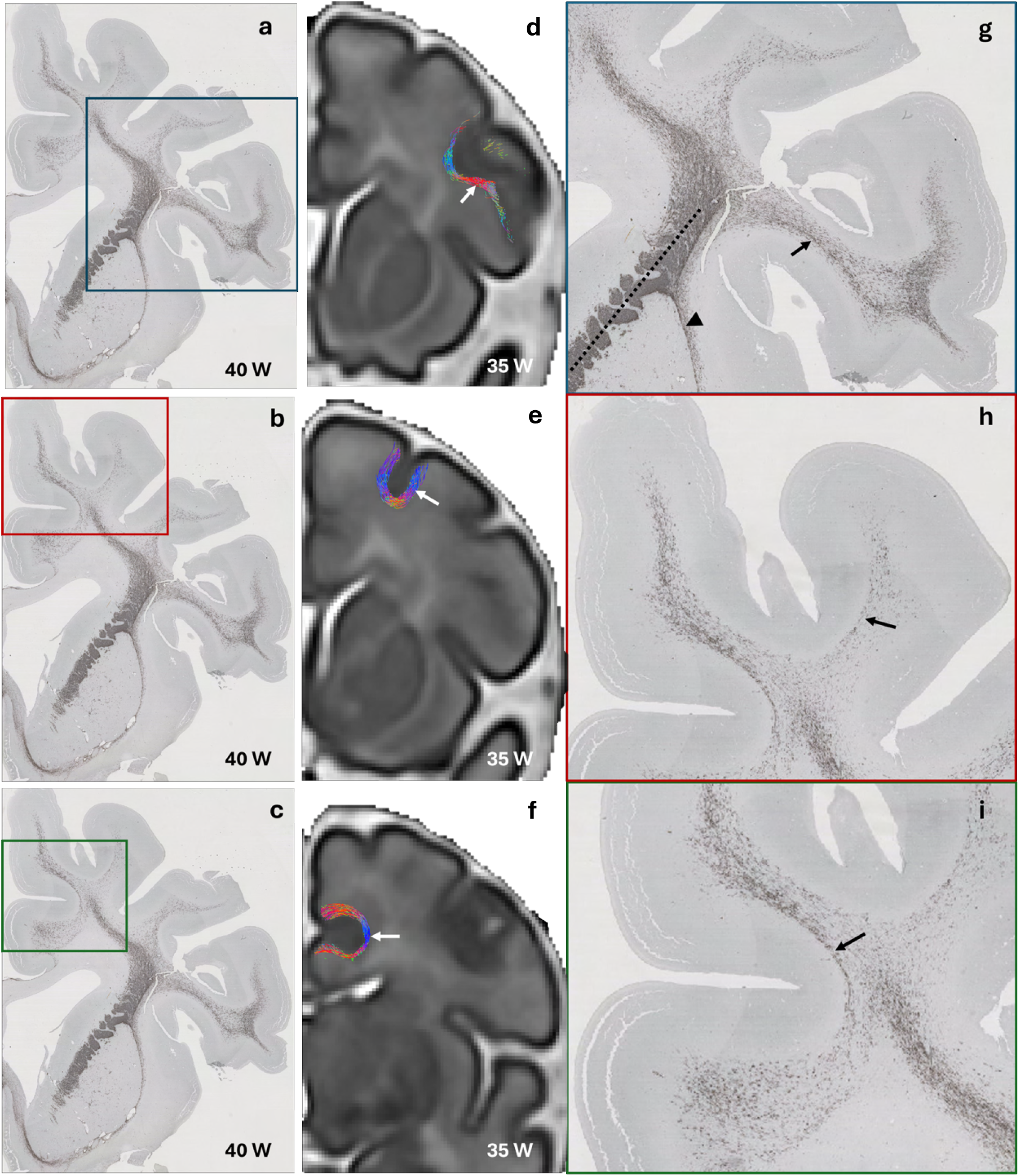
Validation of in-utero MRI-derived short association fibers (SAFs) using ex-vivo Myelin Basic Protein (MBP) staining. (**a, b, c**) MBP-stained coronal sections through the frontal region of a 40 gestational week (GW) human brain reveals multiple axonal fiber systems, including prominent projection fibers passing through the internal (dashed line in **g**) and external capsules (arrowhead in **g**). (**g, h, i**) Higher magnification views of boxed regions in (**a, b, c**) highlight SAFs beneath the cortical layer VI (balck arrows). (**d, e, f** ) Corresponding SAF pathways reconstructed from the dHCP fetal template at 35 GW show narrow, U-shaped pathways (white arrows) whose trajectory and position closely align with those observed histologically. These include connections between the middle frontal gyrus and the opercular portion of the inferior frontal gyrus (**d**), between the superior and middle frontal gyri (**e**), and between the supplementary motor area and the mid-cingulate cortex (**f** ). These findings support the anatomical validity of the proposed in-utero MRI-based analytical framework for capturing early SAF systems and demonstrate close correspondence with histological fiber architecture in the perinatal brain. While histological validation is limited to near-term specimens, the MRI-based reconstructions span a much earlier developmental window, enabling investigation of SAF organization throughout the extremely-preterm, very-preterm, and preterm stages.

### Bundle-level reproducibility reveals spatiotemporal emergence patterns

Building on the successful in-utero reconstruction of SAFs and their confirmed anatomical validity against post-mortem histology, we next examined how these pathways emerge and morphologically change over gestation. Specifically, we assessed the reproducibility of individual SAF bundles across fetuses within each age group to identify robustly expressed pathways and to assess their consistency across age groups, resulting in approximately 250 reproducible bundles per age group. Beyond filtering out spurious streamlines, bundle-wise reproducibility provided a quantitative marker of population-level maturation, indicating when and where coherent SAF geometries begin to consistently manifest across individuals. We defined the temporal onset of each bundle as the earliest gestational age at which it became consistently reproducible across all later groups. Using the 36 GW reproducible set as a reference, we traced each bundle backward to determine when this sustained group-level presence began. To explore regional differences in maturation, bundles were grouped by connected lobes (e.g., Frontal-Frontal, Parietal-Temporal). To examine its associations with cortical developmental timing, bundles were additionally grouped by the folding stage of connected gyri (early, mid, or late) based on established histological timelines (Chi et al., 1977).

The distribution of onset ages revealed pronounced spatial heterogeneity (Figure 4a), with a clear central-to-peripheral maturation gradient. Bundles in parietal and parieto-frontal regions, particularly those associated with the sensorimotor and perisylvian cortices, emerged earliest and remained reproducible throughout the third trimester (Figure 4c). Whereas, bundles in more peripheral regions, including the prefrontal and occipital cortices, emerged later. This spatiotemporal pattern echoes postmortem DTI findings (Takahashi et al., 2012) and parallels the known sequence of cortical maturation (Gogtay et al., 2004; Ducharme et al., 2015; Dubois and Dehaene-Lambertz, 2015; Ducharme et al., 2016). Regionally, early-developing bundles predominantly involved primary sensory and motor areas (e.g., postcentral, supramarginal, and precentral gyri), followed by a progressive anterior expansion from premotor areas to the superior, inferior, and middle frontal gyri, eventually reaching the frontal pole. Posteriorly, development extended from medial occipital areas to the lateral occipital and inferior temporal gyri. This spatial gradient also mirrored the sequence of cortical folding: bundles connecting early-visible gyri tended to exhibit directional coherence and reproducibility at younger gestational ages than those involving mid- or late-developing gyri (Figure 4c). Categorical analyses based on lobar location and folding stage revealed an additional pattern: bundles connecting anatomically or developmentally similar regions (e.g., within-lobe or within-stage) generally emerged earlier than those bridging disparate regions or stages. While the inclusion of intra-gyri bundles may have contributed to the earlier reproducibility within these categories, this alone could not explain the overall trend. The parieto-frontal group was a notable exception, emerging early despite spanning distinct lobes, consistent with the advanced maturation of the frontoparietal system.

**Figure 4.**
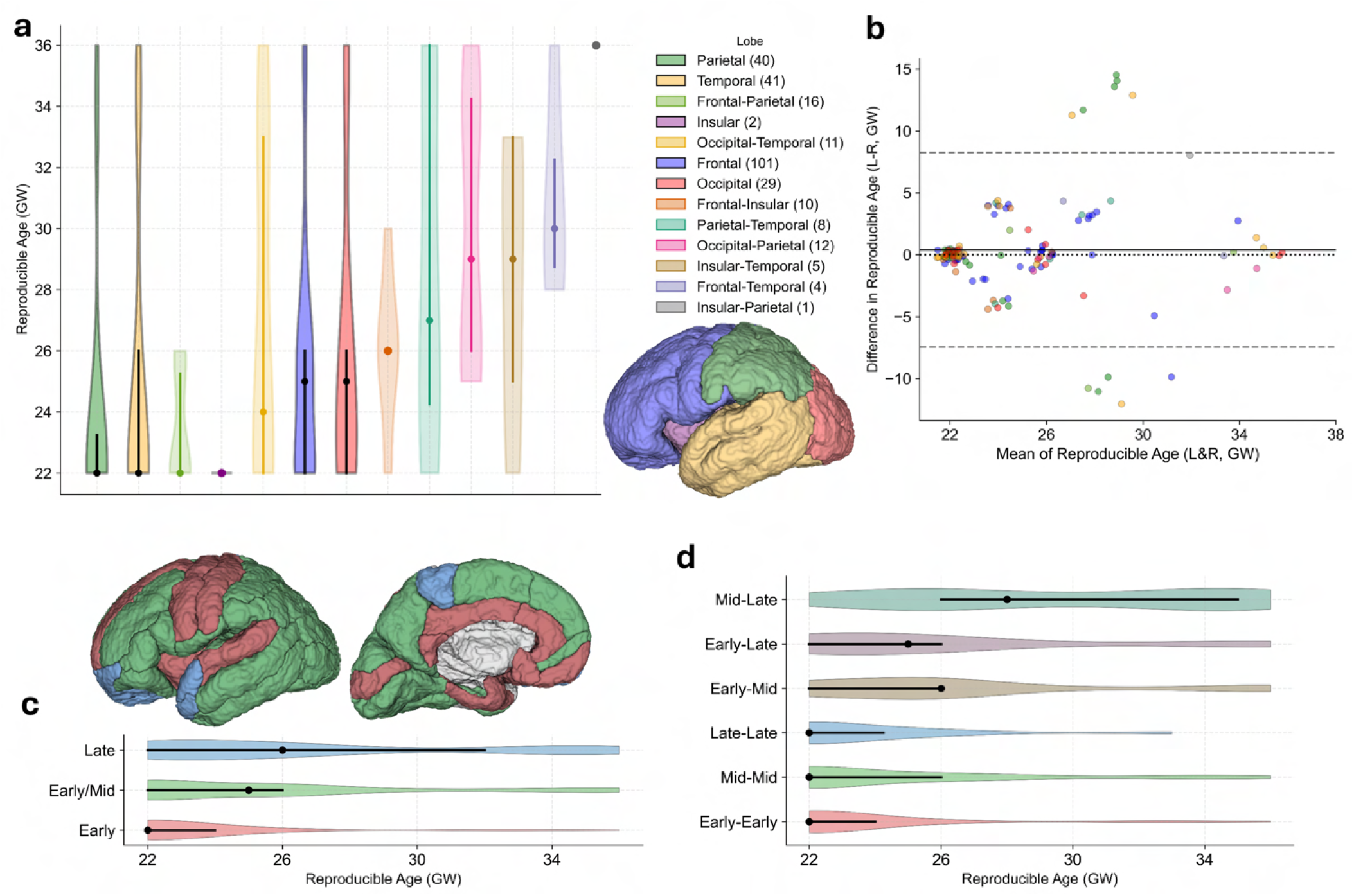
Spatial patterns and hemispheric symmetry of superficial white matter bundle emergence. **a**. Onset of reproducibility grouped by the lobe of the cortical gyri connected by each bundle. Onset is defined as the earliest gestational age at which a bundle could be reliably reconstructed within and across age groups, based on group-wise consistency analysis (see Methods). Bundles are classified by lobar connectivity, with left and right hemisphere bundles considered separately. Violin plots show the distribution of bundle-specific onset ages within each lobar group; central dots represent medians and bars indicate the 25th and 75th percentiles. **b**. Left-right comparison of onset timing among 128 matched bundle pairs. The Bland-Altman plot shows the difference in onset age between left and right bundles against their average. The solid black line denotes the mean difference (0.4 weeks), and the dashed gray lines represent the 95% limits of agreement (mean *±*1.96*×* standard deviation). No significant hemispheric asymmetry was observed (paired t-test, *p* = 0.25), indicating overall symmetry in the anatomical emergence of SAF bundles during fetal development. Eleven bundle pairs fall outside the 95% limits, exhibiting extreme left–right differences. Rather than indicating true hemispheric asymmetry, these likely arise from methodological factors, particularly our conservative, hierarchical emergence criteria, which require consistent group-level presence from the first appearance onward. This criterion may delay confirmation in one hemisphere despite anatomical presence in individual fetuses, due to signal dropout or limited sampling quality within certain age groups, which can hinder group-level consensus despite anatomical presence in individuals. **c**. Onset of reproducibility grouped by the major folding stage of the gyri (Early, Mid, or Late, Table A1) (Chi et al., 1977) connected by each bundle, using a conservative classification: bundles are assigned based on the later-developing gyrus in each pair (e.g., a bundle connecting an early- and a late-stage gyrus is classified as Late). **d**. A finer breakdown by all six gyral pair combinations (e.g., Early-Mid, Mid-Late), capturing intermediate stages of cortical development.

**Figure 5.**
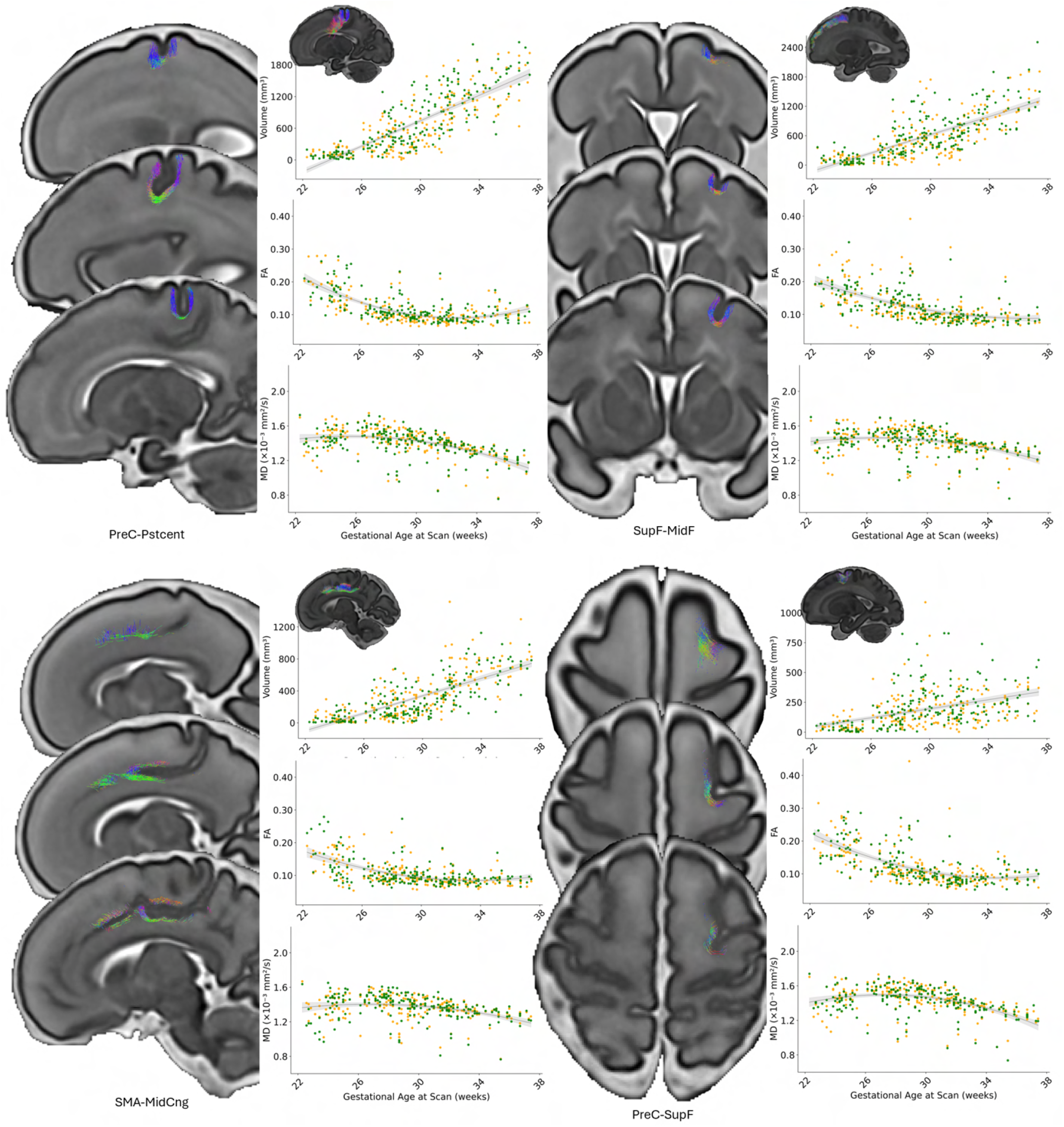
Four SAF bundles in the frontal lobe are shown: PreC-Pstcent (Precentral to Postcentral gyrus), SupF-MidF (Superior to Middle Frontal gyrus), SMA-MidCng (Supplementary Motor Area to Middle Cingulate gyrus), and PreC-SupF (Precentral to Superior Frontal gyrus). For each bundle, developmental trajectories are visualized at 29, 32, and 35 gestational weeks (GW) using fiber pathways intersecting a T2w MRI slice, providing a cross-sectional perspective and highlighting morphological changes relative to the emerging cortical convolutions. Bundle-specific measurements of volume (top; *mm*^3^), free water-eliminated fractional anisotropy (middle; FA), and mean diffusivity (bottom; MD, *×*10^*−*3^*mm*^2^*/s*) are plotted across gestational age. A 3D rendering of the entire fiber bundle at 35 GW is overlaid on a 2D sagittal MRI slice (left view) to show its overall shape and anatomical location, using the same visualization approach as in Figure 1b. FA and MD values represent the median of non-zero values within each bundle mask. Left (orange) and right (green) hemisphere measurements were jointly modeled. A linear model was used for volume, while DTI measures were fit using either linear or quadratic regression based on adjusted *R*^2^. Shaded regions denote the 95% confidence intervals of the fitted trajectories, estimated via nonparametric bootstrapping (1,000 iterations). Rather than forming the static U-shaped configuration commonly seen in adult brains, fetal SAF bundles initially appear as flat structures with 2-3 dominant orientations, and gradually reorganize into tightly bundled, directionally coherent, arcuate pathways as sulci emerge and deepen, as exemplified by the SMA-MidCng pathways. This reshaping follows along subplate-white matter interface in a sensorimotor-to-peripheral gradient, with early curvature observed near the central sulcus and later changes in frontal and occipital poles.

We also assessed hemispheric asymmetry in the onset timing of reproducible bundles by comparing 128 left-right bundle pairs present in both hemispheres. Although a small number of bundle pairs showed notable differences in onset age, the overall difference was not statistically significant (paired t-test, *p* = 0.25). A Bland-Altman analysis confirmed the absence of a systematic trend, with most bundle pairs clustering near the zero-difference line and falling within *±*5 weeks (Figure 4b).

### Trajectories of structural and microstructural maturation in individual bundles

Having identified the spatiotemporal emergence gradient of SAF bundles across the fetal cortex, we next quantified the developmental trajectories of individual bundles, focusing on changes in volume and free-water-eliminated DTI metrics (FA and MD) to capture both macroscopic growth and microstructural maturation. All measurements were computed in each subject’s native space. Volumes were estimated from streamline-defined voxel occupancy, while microstructural measurements were sampled using group-level consensus masks warped to the individual brain.

Figure 5 presents results from four frontal lobe bundles: Precentral-Postcentral (PreC-Pstcent), Superior-Middle Frontal (SupF-MidF), Precentral-Superior Frontal (PreC-SupF), and Supplementary Motor Area-Middle Cingulate (SMA-MidCng). Volumetric expansion was observed across all bundles, though the rate and pattern varied, likely reflecting variations in gyral size, inter-gyral interface extent, and developmental timing. While most bundles exhibited near-linear growth within the studied age range, several bundles showed localized volume surges around 30 GW, such as the PreC-SupF connection. These fluctuations may reflect regional dynamics in subplate expansion and dissolution.

Microstructural maturation followed a more complex, nonlinear trajectory. Free-water-eliminated FA decreased in early gestation, reached a turning point (minimum), and then increased, while MD showed the opposite pattern. The turning points of MD (peaks) were earlier than those of FA, mostly between 26 and 30 GW, likely reflecting maximal subplate expansion, particularly in the frontal associative areas. FA turning points were observed later (after 31 weeks), or in some cases, were not yet reached within the studied age range. Notably, regional variation in microstructural maturation was evident even within the frontal lobe. For example, the PreC-Pstcent bundle showed an earlier FA turning point at 31.5 weeks, compared to later turning points in PreC-SupF (36.6 GW) and SMA-MidCng (32.6 GW), while SupF-MidF exhibited an even later shift that was extrapolated beyond the study range at 39.7 weeks. These diverse trajectories highlight how local connectivity pathways follow distinct maturational schedules even within the same lobe. Such bundle-specific differences likely reflect underlying biological processes. Myelination and axonal organization progress along regionally asynchronous timelines (Flechsig, 1920), while the subplate undergoes dynamic changes in the amount and composition of ECM and changes in its fiber architectonics (Kostović et al., 2014a). Our findings align with histological and imaging literature suggesting that simpler laminar architecture, such as limbic areas, tend to follow simpler growth trajectories, whereas polysensory and high-order association cortices, characterized by greater cytoarchitectonic complexity, exhibit more complex, protracted developmental profiles (Shaw et al., 2008).

Complementary results for additional bundles are presented in the Appendix. These include further frontal pathways (Figure A6; OpIF-OpIF, TriIFG-TriIFG), as well as SAFs in the temporal (Figures A2 and A7; e.g., SupTemp-MidTemp, MidTemp-InfTemp, Ang-MidTemp, Fusiform-InfTemp), parietal (Figures A3 and A8; e.g., Precuneus-PCL, Pstcent-SPL, SPL-IPL, SPL-Precuneus), occipital (Figures A4 and A9; e.g., SupOcc-MidOcc, MidOcc-InfOcc, Calc-Ling, Ling-Fusiform), and insular cortices (Figure A5; e.g., OpIF-Ins, OrbIF-Ins, RolOper-Ins). Full bundle names and abbreviations are listed in Table A1. Together, these analyses provide the first cortex-wide, inter- and intra-gyri characterization of SAF maturation in utero, revealing regionally heterogeneous growth and highlighting the sensitivity of DTI metrics to the underlying developmental processes.

### Spatial heterochrony in SAF maturation revealed by microstructural turning points

Extending from bundle-specific developmental trajectories, we next examined the timing of key microstructural transitions to uncover broader organizational principles of SAF maturation across the cortex. Specifically, we identified turning points in free-water-eliminated FA and MD curves, defined as local extrema in quadratic fits, as temporal landmarks where the direction of tissue property change reverses, indicating a shift in the prevailing biological mechanisms driving maturation. Across the cortex, MD and FA exhibited complementary temporal dynamics (Figure 6). MD turning points consistently preceded those of FA, with MD peaks occurring between 24 and 29 weeks and extending to 31 weeks in the frontal lobe. These early MD peaks likely coincide with transient subplate expansion, a phase marked by loosely arranged axons and increased ECM. The subsequent decline in MD suggests a progressive restriction of water diffusion as axonal organization increases and as the subplate dissolves. FA, in turn, reached its minimum later, generally after 31 weeks, with subsequent increases likely driven by axonal packing, laminar differentiation of the cerebral wall, and early myelination. This temporal offset between MD and FA suggests that each metric captures distinct yet interrelated aspects of SAF development, with MD being more sensitive to early permissive isotropic environment and FA to later-phase fiber organization.

**Figure 6.**
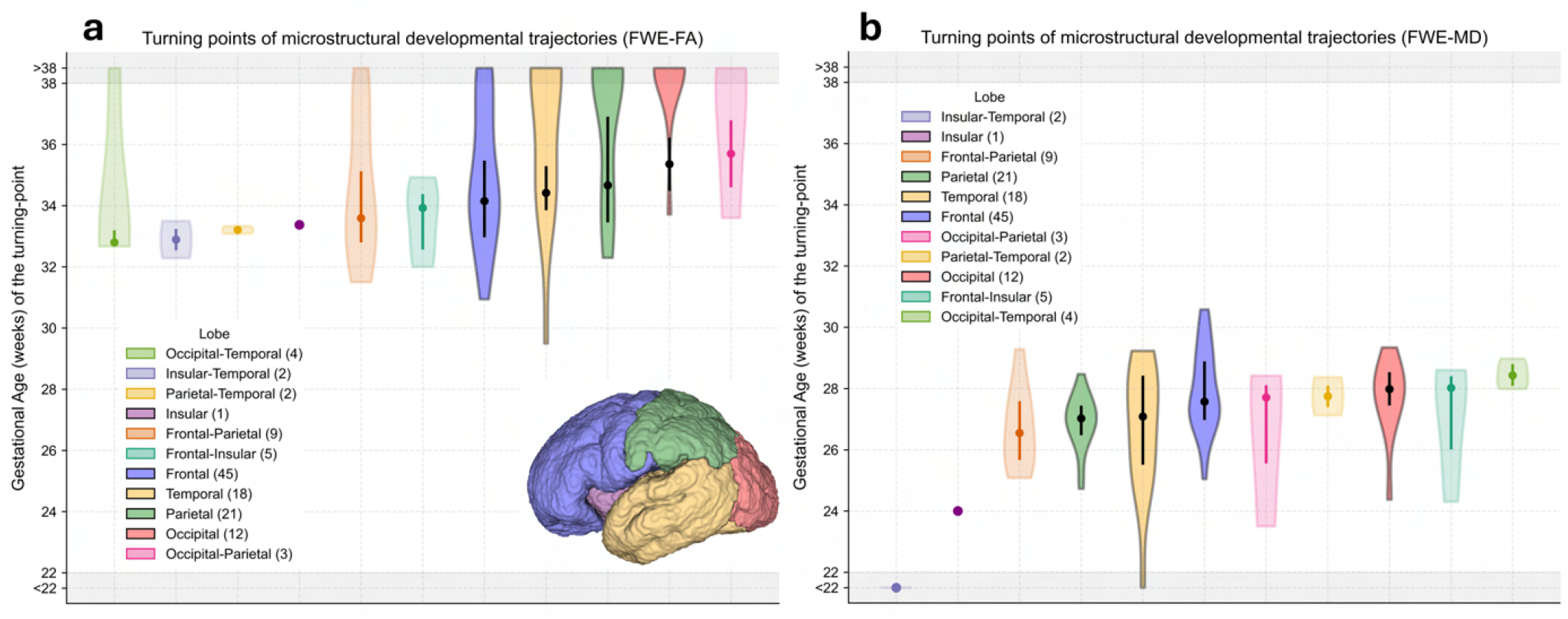
Timing of microstructural maturation of short association fibers (SAFs) across the fetal cortex, assessed by gestational age (weeks) at the turning points of bundle-specific trajectories of free-water-eliminated fractional anisotropy (FWE-FA, **a**) and mean diffusivity (FWE-MD, **b**), grouped by anatomical lobe. Turning points were extracted from the vertex of the best-fitting quadratic model for each bundle and reflect transitional phases in tissue development. Bundles with monotonic trends or with turning-point estimated beyond the observed window (22-38 gestational weeks) were assigned a value of “*>*38 weeks” to indicate that no turning-point was observed before 38 weeks, and were excluded from the statistical summary. Left and right hemisphere measurements were combined for joint modeling and counted once per regression. In all panels, violin plots show the distribution of bundle-specific gestational ages grouped by lobar classification; Summary statistics: central dots for medians and bars for the 25th and 75th percentiles were computed using only values within the observed study range (“22*≤*GA*≤*38 weeks”). The color legend indicates the lobar groups, with the number of contributing bundles shown in parentheses.

**Figure 7.**
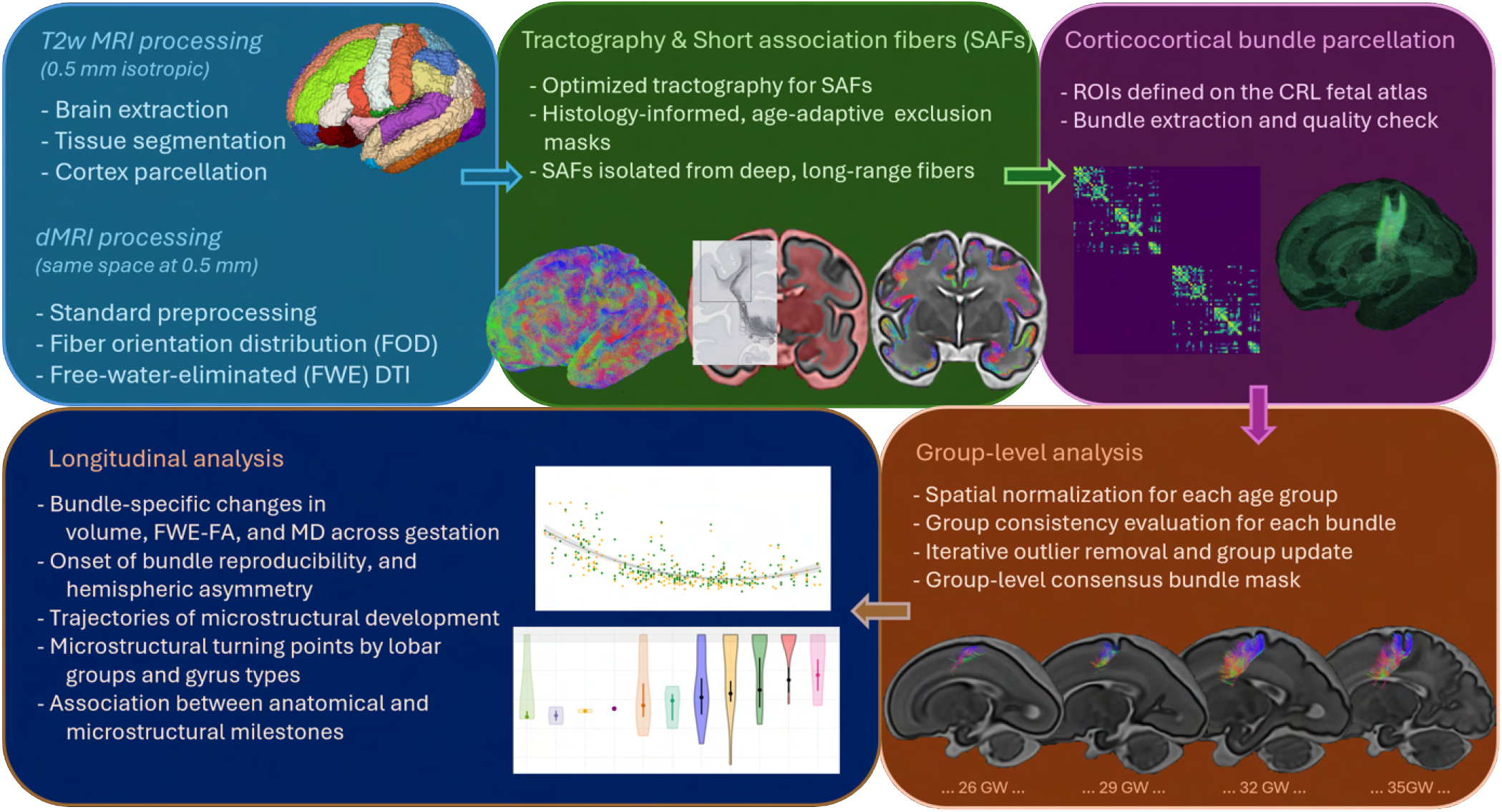
Schematic pipeline of the present spatiotemporal study of short association fiber development in the human fetal brain. The framework integrates gestational age-specific preprocessing, optimized tractography, and histology-informed exclusion mask to isolate SAFs from deep, long-range fibers. Following fiber bundle parcellation, longitudinal analyses characterize bundle-wise trajectories of volume and microstructure (FWE-FA and MD) across gestation. Additional analyses identify the onset of bundle reproducibility, hemispheric asymmetries, and developmental turning points stratified by lobar groups and gyral types. Together, this study represents the first in utero, gyri-level, whole-brain characterization of SAF maturation from 22 to 38 gestational weeks.

**Figure 8.**
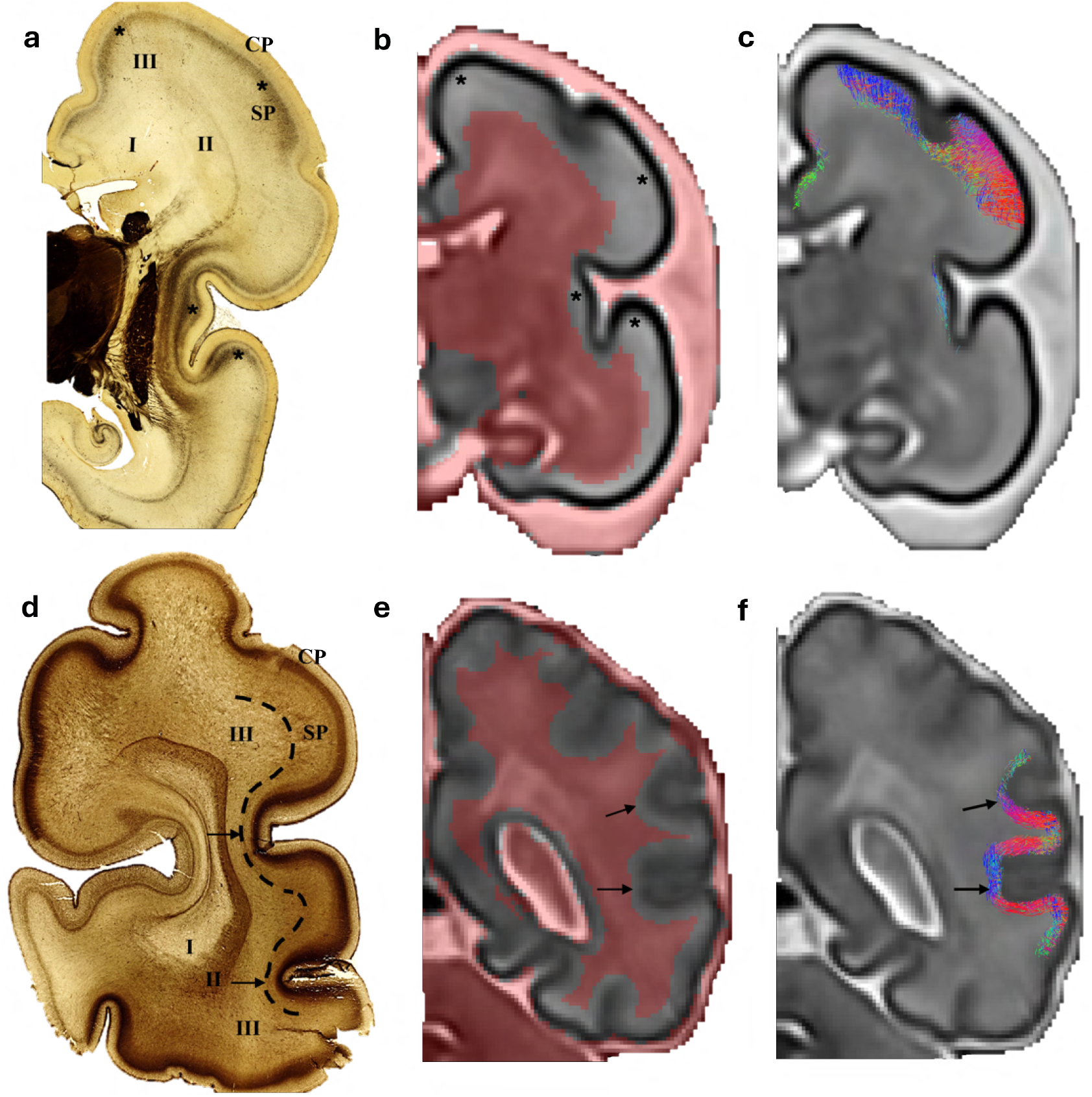
Anatomical localization of the subplate (SP) and the short association fibers (SAFs) presented in this study, as shown by Acetylcholinesterase (AChE) histochemistry (**a**,**d**) and in-utero MR imaging (**b**,**c**,**e**,**f** ) of the fetal brain at very preterm (28 GW; **a-c**) and late preterm (36-37 GW; **d-f** ) stages. (**a**) During early preterm stage, the subplate develops and is the largest compartment of the cerebral wall (Kostović et al., 2014a). The most superficial part of the subplate displays an intense AChE-reactivity and hyper-intensity on T2-weighted MRI (asterisk). The position of the future centrum semiovale (white matter segment III, WMS III) is now largely occupied by the subplate. CP: cortical plate, WMS II: intermediate zone, which consists of sagittal strata and periventricular crossroad of pathways, WMS I: (deep) periventricular zone. (**d**) As brain develops, the subplate diminishes in size, and its former position is progressively occupied by WMS III (centrum semiovale) and newly developing gyral white matter. The subplate dissolution is prominent at the bottom of cortical sulci (arrows). Dashed lines delineate the border between the subplate and underlying centrum semiovale, corresponding to the transition between the superficial and intermediate compartments of the fetal brain. (**b**,**e**) In-utero fiber extraction masks derived from T2-weighted MRI, isolating superficial fibers by retaining only those fully confined to the subplate (or subplate remnant) and gyral white matter. Fibers traversing or terminating in excluded regions (in red), such as the centrum semiovale, subcortical gray matter, or brainstem, are removed. (**c**,**f** ) Reconstructed SAFs, extracted from the filtered superficial tractograms using the corresponding masks in (**b**) and (**e**).

Two large-scale principles of maturation emerged from the distribution of turning points. The first reflects a spatially ordered progression that extends from the central sulcal (fissure of Rolando) toward the lateral convexity (e.g., inferior temporal gyrus) and anterior poles. For example, within the temporal lobe, the Superior-Middle Temporal bundle reached its FA turning point by 36.8 weeks, while the Middle-Inferior Temporal bundle followed later at 40.5 weeks (Figure 4c). A similar pattern was observed in the occipital lobe, where the Superior-Middle Occipital bundle (38.5 weeks) preceded the Middle-Inferior Occipital bundle, which showed a projected FA turning point beyond term (50.5 weeks), suggesting critical microstructural changes after birth (Figure A4). The Lingual-Fusiform bundle, located in the lateral occipitotemporal cortex also followed a protracted trajectory, with a turning point estimated at 42.5 weeks. These spatial patterns mirror known sequences of gyral development (Chi et al., 1977).

Superimposed on this spatial gradient was a second, partially overlapping pattern that followed a functional gradient from primary sensorimotor areas to higher-order association cortices, consistent with the sensorimotor-association (or unimodal-transmodal) axis featured in human cortical organization (Mesulam, 1998; Margulies et al., 2016; Huntenburg et al., 2018; Sydnor et al., 2021). Early-maturing axonal bundles included PreC-PostC (31.5 weeks) and SM-MidCng (32.6 weeks), which are anchored in primary or supplementary motor regions. Whereas bundles associated with higher-level cognitive and executive functions, such as SupF-MidF in the dorsolateral prefrontal cortex, showed later maturation (39.7 weeks). Cuneus-Lingual (37.0 weeks), situated in the primary visual cortex along the calcarine sulcus, reached its turning point earlier than Lingual-Fusiform (42.5 weeks), which lies in the lateral visual association cortex. Within Broca’s region, we observed subtle timing variations: the OpIF-OpIF bundle (34.8 weeks), in pars opercularis (Brodmann Area 44, BA44), exhibited an earlier maturation inflection than TriIFG-TriIFG (36.3 weeks), in pars triangularis (BA45). While modest, this distinction may reflect differential maturation related to their proposed roles in motor aspects of speech versus lexical-semantic processing (Hagoort, 2014). Other variations within Broca’s area, including cytoarchitectonic differences (BA44 dysgranular, BA45 granular) and distinct connectivity profiles (Amunts and Catani, 2015), further scaffold this functional specialization, although specialization in this region remains a subject of ongoing investigation.

Finally, we compared the timing between bundle emergence and microstructural turning points to assess whether anatomical geometry and maturation unfold in synchrony (Appendix Figure A10). While early-emerging bundles often matured earlier, especially in sensorimotor areas, this temporal alignment was not uniformly observed across all bundles. These findings suggest that emergence and maturation, while partially coordinated across regions, are further shaped by region-specific developmental programs.

## 3 Discussion

SAFs form a dense scaffold of local connectivity, facilitating early communication between neighboring gyri during cortical development. In this study, we systematically charted the anatomical and microstructural development of SAFs in the fetal brain, revealing an organized yet heterogeneous maturation process. SAFs were found to emerge early in gestation as loosely arranged, multi-directional pathways and gradually exhibited coherent orientation, and arcuate shapes tightly delineating cortical sulci. This morphological transformation was accompanied by a progressive increase in population-level reproducibility and anatomical specificity. Quantitative analysis of over 250 bundles across the cortex uncovered regionally distinct patterns of formation and maturation, with earlier development in sensorimotor regions and delayed trajectories in higher-order association areas. Microstructural measurements exhibited nonlinear changes over gestation, with turning points in FA and MD revealing spatial heterochrony in tissue maturation. Together, these findings demonstrate that SAF development follows a coordinated yet regionally variable timeline shaped by both spatial and functional hierarchies, providing a foundation for understanding the early emergence and formation of local cortical circuits.

The time of appearance and positioning of SAFs along the radial axis of the cerebral wall in human fetuses can be understood within the broader, sequential yet overlapping maturation of distinct classes of cerebral pathways.

Our findings refine this understanding by charting the in-utero trajectory of SAF maturation with unprecedented spatiotemporal detail, revealing that SAFs emerge during mid-to-late phase of cortical connectivity development, after long-range projection pathways are in place, but before intracortical circuits develop postnatally. This temporally orchestrated sequence begins with modulatory afferents from brainstem nuclei, arriving before 8 GW and targeting the marginal zone and subplate. Shortly thereafter, afferents from the basal forebrain and other subcortical telencephalic nuclei begin to populate the subplate. Following the establishment of the CP around 8 GW, thalamocortical projections undergo extensive development, with fibers penetrating the CP only much later, around 22-23 GW, after forming transient synapses in the subplate. The corpus callosum begins to form after 10 GW, likely arising from early layer III pyramidal neurons whose axons project contralaterally. Around this period, one of the earliest short corticocortical circuits, the reciprocal connections between the entorhinal cortex and hippocampus (archicortex), also begins to form, supporting early limbic network development before the maturation of neocortical associative systems (Hevner and Kinney, 1996). Long-range associative fibers, also originating from layer III pyramidal neurons, begin to emerge at preterm stages (subsequent to callosal formation), and exhibit both ipsilateral and contralateral collaterals. The uncinate fasciculus is among the earliest long association tracts to appear (Huang et al., 2009), and by approximately 20 GW, most major pathways become identifiable in dissection studies (Horgos et al., 2020). The SAFs described in the present study, develop in parallel with these long-range systems but are more closely linked to the expansion of local cortical circuits (Reveley et al., 2015). Finally, intracortical fibers, including tangential connections within the cortical layers, predominantly develop after birth, as supported by postnatal histological evidence (Burkhalter et al., 1993).

The fetal-stage development of SAFs coincides with a critical transitional window of cortical network assembly, as local circuits begin to form on the scaffold of long-range connectivity. Anatomically, SAFs are confined to the superficial compartment between the CP and deeper WM segments, a region dominated during this period by the subplate. Their development, therefore, closely parallels subplate maturation, which presents a major hub of early synaptogenesis, axonal pathfinding, neuron-neuron-glia interaction, and early network coordination (Kostović et al., 2021). Notably, SAF expansion accelerates between 28 and 32 GW, during the peak thickness of the subplate and a period of increased vulnerability to perinatal brain injury. These spatiotemporal patterns suggest that SAFs help establish local communication within the global brain network and may serve as in-utero markers of cortical circuit maturation and early neurodevelopmental risk.

Spatial heterochrony across structural and microstructural milestones emerged as a defining feature of SAF development, with anatomical establishment and underlying biological processes following regionally asynchronous timelines. Bundles in primary sensorimotor regions appeared earliest, including parietal and frontal-parietal areas, e.g., the prominent precentral-postcentral U-fiber system which has been well-described in the mature brain (Zhang et al., 2010; Oishi et al., 2011; Catani et al., 2012; Guevara et al., 2017; Magro et al., 2012; Román et al., 2017; Rojkova et al., 2016; Pron et al., 2021).

The spatiotemporal gradient of SAF development was coordinated with the broader principles of fetal brain maturation, in which regional specialization, laminar organization, gyrification, and subplate dynamics, influence and are influenced by the maturation of cortical connections (Essen, 1997; Kostović et al., 2021). Early structural coherence in primary sensorimotor bundles aligns with functional observations in preterm neonates: coordinated motor and sensory bursts are detectable by EEG as early as 26 GW (Vanhatalo and Kaila, 2006). Complementary animal studies show that spontaneous retinal activity propagates across the visual system before sensory experience begins (i.e., before eye opening), suggesting that coherent structural pathways can precede and help organize early functional activity (Ackman et al., 2012). Within this study, our observation of early and robust SAF development in the parietal lobe (postcentral gyrus; superior/inferior parietal lobules; precuneus) corresponds with a marked shift in sensorimotor integration and motor coordination in fetal behavior (Vasung et al., 2023). In addition, tactile input begins to be coordinated with motor output, reflecting increased engagement of the parietal and frontal networks and emerging multi-sensory integration, which parallels the early maturation we observed for frontal-parietal connections. This also includes the early coordination of auditory input with motor activity, likely mediated by a network involving the posterior parietal cortex, superior temporal gyrus, inferior frontal gyrus, and angular gyrus (Catani et al., 2017), and includes the emergence of resting-state networks (including auditory, somatosensory, motor, and frontoparietal) during the second and third trimesters (Doria et al., 2010; Jakab et al., 2014). Together, these converging lines suggest that early-developing local association pathways may contribute to the scaffolding of functional circuits during a critical window of network formation.

The region-specific nonlinearity of FA and MD trajectories and the systematic lead of MD peaks over FA minima support a multiphasic model of SAF maturation: early phases dominated by subplate expansion with a large and hydrated extracellular space (rising MD, declining FA); a transitional phase characterized by the accumulation of loose plexiform network of fibers and post-migratory cells (declining MD); and later phases marked by subplate ECM dissolution, emergence of gyral WM, tightly packed and orderly arranged axons, and myelination that strengthen directional coherence (rising FA), with earlier timing in primary sensory and motor regions. Interestingly, the transitional phase characterized here, manifesting as a decline in MD prior to FA increase, largely corresponds to the “pre-myelination” stage described in earlier imaging studies, which noted restricted water diffusion at developmental stages proceeding histologically confirmed myelination (Hüppi et al., 1998a,b; Wimberger et al., 1995). This phase is thought to be associated with increases in axonal caliber, the initial ensheathment of axons by oligodendrocytes, and, within the subplate zone, a 3D reorganization of the axonal plexus (Inder and Huppi, 2000; Kostović et al., 2002; Cowan et al., 1984). Although free-water-eliminated DTI mitigates contamination from extracellular fluid, the resulting measures still reflect a composite change of tissue microstructure.

The timing of anatomical convergence varies across bundles, and it does not always parallel microstructural shifts, suggesting early geometrical consolidation followed by protracted tissue-level refinement shaped by local factors (Vasung et al., 2016). The decoupled timing echoes observations from infancy that early myelination onset is not invariably followed by early myelin maturation (Kinney et al., 1988). Region-specific or multivariate modeling, akin to work on deep WM bundles (Zanin et al., 2011), may help identify organizing principles that govern how macroscopic geometry and microstructure advance together in some regions and proceed on offset schedules in others.

Placed alongside existing preterm models, our in-utero findings show both shared patterns and important divergences that arise from differences in developmental coverage (a large cohort spanning 22-38 GW), biological milieu (in utero vs. ex utero), and anatomical specificity. Prior “prenatal” SWM inferences predominantly come from neuropathological studies obtained postmortem, or very preterm neonates (*<*30 GW) scanned ex utero. Those studies describe spatially heterogeneous maturation, with faster FA increases in primary sensorimotor cortex than in association areas (Smyser et al., 2016; Yuan et al., 2023), a pattern that parallels our regional gradients. Key differences, however, highlight why an in-utero baseline is needed. First, most preterm studies report linear FA increases from approximately 28-32 GW onward, consistent with their sampling window, whereas our in-utero coverage back to 22 GW and bundle-level analysis reveal earlier, bundle-specific nonlinear phases that precede later increases. These early phases, including an initial FA decline, would be difficult to detect in cohorts that begin after 26 GW and include few observations between 26-30 weeks, limiting the power to identify earlier stages of the development, particularly those occurring before cerebral pathways are established. Second, in preterm cohorts, brain injury can obscure developmental patterns due to anterograde and retrograde degeneration, and compensatory reorganization (Goldman and Alexander, 1977). Axonal pathways are still growing in the early preterm stage, making WM in preterm infants significantly different from that of term newborns (Kostović et al., 2019; Volpe, 2009; Counsell et al., 2003; Batalle et al., 2017). Moreover, the developmental milieu differs fundamentally: very preterm infants mature ex utero, where altered sensory input, stress, and nutrition can influence neurodevelopmental trajectories, sometimes delaying features (e.g., sulcation (Barkovich, 2005), widespread FA reductions (Anjari et al., 2007)), and sometimes accelerating others (e.g., experience-driven FA increases in visual cortex (Ment et al., 2009)). A third difference lies in anatomical precision. Many prenatal studies approximate SWM, WM (Schneider et al., 2016, 2007, 2009; Partridge et al., 2004; Calixto et al., 2025a), or the subplate (Gupta et al., 2005), using fixed-depth sampling beneath the cortex or manually placed ROIs, risking inclusion of adjacent compartments, and overlooking regional and gestational dynamics of the fetal cerebral wall. In contrast, our framework leverages refined spatial grids (0.5 mm isotropic), anatomically guided tractography, bundle-specific modeling, and gestational age-adapted tissue segmentation to explicitly account for the developing interface between superficial and deeper compartments, enabling gyri-level resolved SAF mapping across the fetal cortex. Together, these features underscore the unique value of in-utero diffusion MRI in establishing a normative reference for SAF development, free from prematurity-related confounds and optimized for fetal neuroanatomy.

MRI-histology correspondence supports the anatomical specificity of the in-utero SAF reconstructions, strengthens the interpretability of diffusion-derived measurements as highly sensitive in-vivo indices of tissue changes within the appropriate laminar context, and highlights the advantages of in-utero diffusion MRI, i.e., fine temporal sampling, 3D quantification and visualization, and whole-brain coverage, relative to the necessarily sparse sampling of postmortem sections. More broadly, these results illustrate the value of anatomically grounded in-utero imaging for resolving early SAF organization during its emergence, and provide a scaffold for future integrations with invitro postmortem imaging and molecular datasets to probe regional vulnerability and the development of circuit specialization.

Beyond establishing an in-utero baseline and aligning MRI with histology, we introduce a set of methodological advances that are critically needed for tracking and visualizing connectivity in the developing human brain (Ouyang et al., 2017). First, we operationalize an anatomically faithful definition of fetal SAFs, confined to the subplate and emerging gyral WM, using a histology-informed, “inside-out” peeling strategy and age-adaptive tissue segmentation that respects the transient subplate zone and the developing SP–WM interface. Second, we pair anatomically constrained, ensemble tractography with gyri-resolved parcellation and a hierarchical quality-control workflow (individual pruning → group normalization to weekly → templates bundle-wise reproducibility within and across age groups). This supports gyri-level mapping of short-range pathways across 22–38 GW, enabling bundle-wise developmental profiling and spatiotemporal modeling of both global maturation patterns and short-range structural connectivity across the fetal cortex. Third, we quantify maturation with bundle-specific macrostructural features and free-water-eliminated DTI metrics, which help mitigate fluid contamination in the highly hydrated fetal brain.

Despite these strengths, several limitations frame interpretation and naturally suggest next steps. (i) Developmental fate. Within the fetal window, we cannot determine the extent to which early SAFs confined to the superficial compartment eventually contribute to long-range associative systems, which are progressively displaced into deeper WM with maturation. (ii) Temporal sampling. Our cross-sectional design (22–38 GW) precludes within-subject developmental trajectories and leaves some turning points extrapolated beyond term. Extending future analyses into immediate postnatal windows would refine inflection ages. (iii) Microstructural modeling. Free-water–eliminated DTI improves specificity and is easy to interpret, but DTI remains a simplified model of complex microstructure. Adding multi-compartment modeling and higher-*b* shells, where feasible, should sharpen biological attribution (e.g., axonal density versus anisotropy). (iv) Group-level masking. Consensus bundle masks enhance robustness for regression yet can attenuate subtle individual variations, a trade-off inherent to large-scale, automated pipelines. (v) Conservative estimation. Our estimates of SAF onset timing are deliberately conservative, anchored to persistent group-level presence across all subsequent age groups. This approach strengthens robustness and prevents from spurious detections, but may lag behind actual anatomical emergence. Similar delays have been noted in prior fetal MRI studies, where sulcal appearance lags behind postmortem reports (Levine and Barnes, 1999) and initial identification occurs before consistent presence (Garel et al., 2001; Glenn and Barkovich, 2006). This conservative criterion may exaggerate left–right differences when emergence is confirmed in one hemisphere but not the other due to signal dropout or small group sizes with variable data quality, where our group-level criteria are harder to meet. Accordingly, extreme hemispheric differences (over 10 weeks in some cases) likely reflect methodological constraints rather than genuine asymmetry.

Looking ahead, several opportunities could broaden the scientific reach of this work. Region-focused analyses (e.g., perisylvian language cortex) can test for subtle asymmetries and specialized timelines that whole-brain summaries may dilute. Multimodal extensions pairing fetal fMRI with bundle-wise diffusion metrics would probe evolving structure–function coupling in utero. Applying the pipeline to high-risk cohorts or fetuses with high risk genotypes will help contextualize deviations against the normative reference established here and refine early biomarkers. Finally, community release of code and quality-controlled results will enable benchmarking, encourage harmonization efforts, and test-retest studies, while integrations with histology and transcriptomic dataset can link regional trajectories to underlying cellular programs, improving mechanistic interpretations of SAF maturation.

## 4 Methods

We conducted the study in five main steps using fully automated methods (Figure 7), including 1) data inventory and processing, 2) fiber tracking and isolating superficial fibers from deep, long-range fibers, 3) parcellation of inter- and intra-gyri short association fiber (SAF) bundles, 4) group-wise evaluations for each age group, and 5) longitudinal modeling of bundle-specific morphological and microstructural changes across gestational ages, and lobe-wise analysis. The first three steps were progressively optimized through manual inspections and comparison with neuroanatomy knowledge.

### Datasets and MRI processing

The brain specimens used for postmortem histological comparison were obtained from the Zagreb Collection of Human Brains (Kostovic et al., 1991), a well-established and diverse repository of human fetal brain tissue. The histology study was conducted according to the guidelines of the Declaration of Helsinki, and approved by the Ethics Committee of the School of Medicine University of Zagreb (protocol number Ur. Broj: 251-59-10106-23-111/158, Klasa: 641-01/23-02/01).

Fetal brain MR imaging from the Developing Human Connectome Project (dHCP^2^) was utilized for the study of SAFs in utero. This dataset presents the highest quality open-source data for fetal brain MRI, allowing for cutting-edge and reproducible studies by the entire scientific community. Diffusion-weighted MRI (dMRI) data was collected with a combined spin echo and field echo sequence at 2 mm isotropic resolution (Hutter et al., 2018), using a multi-shell diffusion encoding that consists of 15 volumes at b=0 *s/mm*^2^, 46 volumes at b= 400 *s/mm*^2^, and 80 volumes at b= 1,000 *s/mm*^2^ (Tournier et al., 2020).

Standard preprocessing operations that have been performed on the released data include denoising, eddy current and susceptibility-induced distortion correction, and motion correction (Bastiani et al., 2019; Christiaens et al., 2021). We linearly up-sampled the data to an isotropic voxel size of 0.5 mm. The MRI data and intermediate results derived from them were standardized to ‘Right-Anterior-Superior’ (RAS) orientation. Detailed cortex parcellation, and fetal brain tissue segmentation including transient zones such as the subplate, were obtained by non-linearly aligning the T2-weighted MRI and propagating the CRL fetal brain atlas^3^ (Gholipour et al., 2017) of matched age to individual space (Avants et al., 2009).

The classical model for estimating the local orientation of prominent fibers in each imaging voxel is DTI. Although DTI is inherently limited to representing a single fiber orientation per voxel, it remains a widely used and robust approach, particularly in situations where more complex models cannot be reliably applied. Methods such as spherical deconvolution estimate a full fiber orientation distribution (FOD) that has the potential to resolve crossing fibers. These methods, however, require more measurements and higher signal to noise ratio for reliable model fitting. Moreover, recent studies have shown that standard FOD estimation methods are not be optimal for early brain development, where the tissue microstructure is vastly different from adult brains (Pietsch et al., 2019; Kebiri et al., 2024). In this work, we adopted a practical strategy to choosing the suitable model of local fiber orientations by computing and assessing four different representations derived from the dMRI data. (1) Single-shell single-tissue constrained spherical deconvolution (CSD) using the *b* = 0 and *b* = 1, 000*s/mm*^2^ data (Tournier et al., 2007); (2) Multi-shell multi-tissue CSD using the data in all three shells (Jeurissen et al., 2014); (3) DTI estimated with a weighted linear least squares method; (4) Sharpened diffusion orientation distribution function (dODF) computed from the diffusion tensor (Descoteaux, 2008; Aganj et al., 2010). We computed tractography using these four inputs models, followed by visual inspection and qualitative comparison of the resulting whole-brain tractograms and the extracted superficial bundles. Based on this assessment, we chose the DTI-derived dODF for subsequent analysis, as it provided the most extensive coverage of superficial fibers within gyri compared to the alternatives. Finally, we computed diffusion tensor scalar measures, i.e., fractional anisotropy (FA) and mean diffusivity (MD), from both the standard DTI and free water-eliminated DTI (Pasternak et al., 2009).

### Tractography and SAF systems

To seed and constrain fiber tracking using anatomical priors, we computed five-tissue-type (5TT) segmentation maps from the tissue segmentation provided in the dHCP dataset (Makropoulos et al., 2014), as transient zones in the fetal brain did not have to be distinguished for this step. The 5TT map is a 4D image where the last dimension comprises five mutually exclusive channels: cortical gray matter, subcortical gray matter, white matter, cerebrospinal fluid (CSF), and pathological tissue (which did not exist for the normal fetal brains used in this work). We constructed the 5TT maps by preserving original cortical GM and WM labels, inserting an empty lesion channel, reassigning the brain stem and hippocampus to the subcortical GM channel, and including the lateral ventricles in the CSF channel.

Because the original tissue segmentation tended to overestimate cortical GM, often extending into the subplate and causing “closed gyral crowns”, we refined the segmentation to restore biologically accurate boundaries. These corrections were critical to allow streamlines traverse the expected anatomical zones such as the gyral crown and external capsule. In details, we dilated the white matter mask by one voxel (∼0.5 mm), identified its overlap with the cortical GM label, and reassigned the overlapping regions to white matter. A similar correction was applied to the subcortical GM label, particularly to restore the intervening white matter structures in the insular region.

Given the variability in the size and shape of different superficial fiber bundles, a single tractography setting was not adequate to reconstruct all bundles. Therefore, we adopted an ensemble tractography strategy, where we used a set of 10 different parameter configurations to compute the tractography. Each setting in the ensemble used different values for the angular threshold between successive propagation steps (70-90 degrees), FOD amplitude cutoff to terminate the tracking (10^*−*9^-10^*−*3^), and FOD power (3-5). The ensemble design was intended to cover a wide range of tractography settings to enable a more complete coverage of SWM. All tractograms were generated using a probabilistic tracking algorithm (iFOD2) (Tournier et al., 2010) with the sharpened DTI-derived dODF as the input. The minimum and maximum fiber lengths were set at 4 mm and 45 mm, respectively. For each of the 10 parameter configurations in the ensemble, 100,000 streamlines were generated. Therefore, the final ensemble tractogram obtained by merging the 10 ensemble results, consisted of one million streamlines.

To isolate the superficial fiber system, we applied a biologically informed exclusion strategy. We first removed streamlines that passed through or terminated in subcortical gray matter, brain stem, or cerebellum (to exclude projection pathways). Deep white matter fibers were then excluded using a developmental stage-specific strategy based on the presence of transient zones (to exclude long association pathways). For fetuses younger than 32 weeks, deep white matter was defined as the intermediate zone; only streamlines confined within the subplate were retained. The intermediate zone mask was eroded by one voxel (∼0.5 mm) to mitigate registration and partial volume errors. For fetuses older than 32 weeks, where the subplate is no longer separately labeled, deep white matter was defined as the eroded white matter core (two iterations, around 1-2 mm below the CP). This age-adaptive exclusion strategy ensured consistent delineation of the superficial system across developmental stages.

### Gyri-level SAF bundle parcellation and reproducibility assessment

Following the extraction of SAFs constrained to the superficial compartment, we applied a hierarchical evaluation procedure, as summarized in Algorithm A1. At the individual level, candidate bundles were extracted from the subject-specific tractograms using a parcellation-driven approach based on a gestational age-matched fetal cortical atlas (Gholipour et al., 2017). Streamlines were grouped by the gyri they connect (Figure 1d), yielding approximately 300 intra- and inter-gyri candidate bundles per brain. The full list of gyri used to define candidate connections is provided in Appendix Table A1. Pairwise bundle connectivity at 36 GW is visualized in Figure A11.

At the individual level, we investigated an outlier removal method that scores each streamline in a bundle by its similarity to the other streamlines in the bundle, i.e., cluster confidence index (Jordan et al., 2018). Streamlines falling below the 3rd or 5th percentile of the bundle-specific score distribution were flagged as potentially spurious and removed. We validated this pruning strategy with the GA35 group to be consistent with the histological validations. Whereas visual inspection confirmed that pruning effectively eliminated irregular trajectories in subjects with low image quality, in most subjects the initial unpruned bundles already showed coherent trajectories; pruning in these cases primarily removed peripheral streamlines at the edges of tube-like bundles. To preserve the full extent and anatomical shape of each connection, we ultimately retained the original, unpruned bundles for all subjects in downstream analyses. Bundles with fewer than 100 streamlines were excluded to ensure robustness in later group-level evaluation.

To assess reproducibility across subjects, we performed nonlinear registration of each subject’s diffusion space to their age-matched template in an atlas reconstructed from the dHCP dataset (Uus et al., 2023). In this common space, we computed the binary mask for each bundle by thresholding its streamline density map to retain voxels traversed by at least the 5th percentile of streamlines. Pairwise Dice coefficients were then calculated between each fetus’s bundle mask and the group-consensus bundle mask that was obtained by arithmetic averaging.

To identify bundles with consistent anatomical presence across subjects, a two-step quality control procedure was applied. First, individual bundles were retained if their Dice coefficient of the overlap with the group-averaged bundle mask exceeded a minimum threshold (Dice ≥ 0.1). This liberal threshold was chosen solely to exclude invalid cases (e.g., empty or NaN masks) rather than to assess anatomical precision. Second, a bundle was considered group-wise valid if more than half of the subjects in the group exhibited a valid instance. This criterion ensured that retained bundles reflected consistent, non-random anatomical features rather than subject-specific artifacts. Following this initial filtering, we further refined the group composition by performing iterative subject-level outlier removal. This step aimed to exclude subjects whose bundles globally deviate from the rest of the group, thereby improving the overall group coherence. In each iteration, the subject whose exclusion most improved the mean Dice coefficient across bundles was removed. This process continued until improvements were no longer statistically significant (p *>* 0.01), negligible (improvement in average Dice coefficient 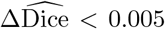), or the group size fell below five subjects.

Using the final retained subject set for each age group, we recomputed group-consensus bundle masks and identified the reproducible short association connections. In each age group, approximately 250 bundles passed quality control and were retained for bundle-level microstructural evaluation and developmental milestone analysis.

To facilitate targeted analysis and visualization, we selected a subset of SAF bundles that connect major gyri for detailed evaluation based on their developmental relevance and consistency across subjects. Specifically, we included the connection between the precentral and postcentral gyri, which are from the primary region of cerebral development and among the earliest-developing short association pathways. Additional regions of interest were defined around early-forming sulci, such as the lateral sulcus and calcarine sulcus, which are typically identifiable by 23-24 weeks of gestation. Also, these bundles have been reported for high reproducibility in prior SAF studies on adults (Zhang et al., 2010; Magro et al., 2012; Van Dyken et al., 2024). These regions were mapped onto the cortical parcellation atlas and used to extract and visualize representative bundles from our tractography results across gestational age groups.

### Bundle-specific longitudinal analysis

Free water-eliminated DTI was computed using a two-compartment model consisting of an isotropic compartment to represent the free water and an anisotropic compartment modeled with a standard diffusion tensor (Pasternak et al., 2009). Fractional anisotropy (FA) and mean diffusivity (MD) were computed from the diffusion tensor compartment.

All quantitative measures were computed in each subject’s native space. Bundle volumes were estimated by counting the number of unique voxels traversed by at least one streamline and multiplying by the voxel size. DTI measures, including FA and MD (or their free-water corrected counterparts), were summarized as the median value within group-averaged bundle masks. Gestational age was treated as a continuous floating-point variable to preserve temporal resolution.

Regression modeling of developmental trajectories was implemented as follows. For each bundle, linear and quadratic models were fit to the observed age-related data. Model selection was based on the adjusted coefficient of determination 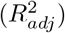, which penalizes model complexity. Subsequently, the selected model and nonparametric bootstrapping (1000 iterations) were used to estimate uncertainty bounds. In each iteration, a sample of the same size as the original dataset was taken from the dataset with replacement, and the selected model was fit to the bootstrap sample. This procedure resulted in 1000 regression coefficients and predictions. At each gestational age, the 95% confidence interval was computed as the 2.5^th^ and 97.5^th^ percentiles of the bootstrapped predictions. The resulting bounds characterize the uncertainty in the estimated mean trajectory, rather than the variability of individual observations.

For bundles that were successfully reconstructed in both hemispheres, bilateral data were combined for regression but plotted using distinct colors to preserve lateralization information.

## Acknowledgment

This research was supported in part by the National Institute of Neurological Disorders and Stroke under award number R01NS128281; the Eunice Kennedy Shriver National Institute of Child Health and Human Development under award number R01HD110772; and by the Croatian Science Foundation grant HRZZ-IP-2022-10-5975 (ŽK) and HRZZ-IP-2024-05-7157 (IK). The content of this publication is solely the responsibility of the authors and does not necessarily represent the official views of the NIH.

Data were provided by the developing Human Connectome Project, KCL-Imperial-Oxford Consortium funded by the European Research Council under the European Union Seventh Framework Programme (FP/2007-2013) / ERC Grant Agreement no. [319456]. We are grateful to the families who generously supported this trial.

## Code and Data Availability

The fetal-specific diffusion MRI analysis pipeline developed in this study will be made publicly available at Gitlab (https://gitlab.com/blibli/fetalSAF). This includes code for image preprocessing, age-adaptive tissue segmentation, tractography, short association fibers extraction, and bundle-wise modeling. In addition, curated outputs derived from the fetal MRI data, such as tract density maps, group-consensus bundle masks, and spatiotemporal summary measures, will be shared via an open-access repository [e.g., Zenodo] to facilitate reuse and further analysis. Histological reference materials are not included. The dHCP imaging data used in this work is publicly available via the National Institute of Mental Health Data Archive (NDA, https://nda.nih.gov/).

## Footnotes

1 https://www.developingconnectome.org/project/

2 https://www.developingconnectome.org/project/

3 http://www.crl.med.harvard.edu/research/fetal_brain_atlas/

## Notes

### Competing Interest Statement

The authors have declared no competing interest.

