## Appendix Table 1 and figures for "Insights into Temporal and Spatial Dynamics of Short Association Fiber Formation in the Human Fetal Brain"

#### Trajectories of structural and microstructural maturation in individual bundles

Full bundle names and abbreviations are listed in Table A1.

Complementary results for additional bundles are presented in the Appendix. These include further frontal pathways (Figure A5; OpIF-OpIF, TriIFG-TriIFG), as well as SAFs in the temporal (Figures A1 and A6; e.g., SupTemp-MidTemp, MidTemp-InfTemp, Ang-MidTemp, Fusiform-InfTemp), parietal (Figures A2 and A7; e.g., Precuneus-PCL, Pstcent-SPL, SPL-IPL, SPL-Precuneus), occipital (Figures A3 and A8; e.g., SupOcc-MidOcc, MidOcc-InfOcc, Calc-Ling, Ling-Fusiform), and insular cortices (Figure A4; e.g., OpIF-Ins, OrbIF-Ins, RolOper-Ins).

#### Cross-analysis of anatomical emergence and microstructural maturation

To evaluate whether the anatomical emergence of SAF bundles is temporally aligned with their microstructural maturation, we performed a cross-analysis of two developmental milestones: the age at which bundles became anatomically reproducible (i.e., showed consistent spatial organization across individuals), and the turning points of diffusion metrics (FA and MD), which indicate local extrema in microstructural trajectories. While these metrics capture distinct developmental aspects, they may jointly inform the sequence of SAF maturation.

Visual inspection of the scatter plots (Figure A9) revealed a lack of temporal coupling between the anatomical and microstructural milestones. For example, bundles with earlier anatomical coherence did not necessarily exhibit earlier FA or MD turning points. Similarly, bundles with later MD turning points (possibly reflecting prolonged subplate development) did not always show correspondingly delayed FA turning points, which may instead relate to different underlying processes such as subplate dissolution, axonal packing or myelination.

These observations were supported by linear regression analysis. The slope was not statistically significant for either FA ($p = 0.418$, $R^2 = 0.0028$) or MD ($p = 0.968$, $R^2 < 0.0001$), indicating no significant linear relationship between anatomical onset and diffusion-based maturation across bundles. Instead of a globally uniform temporal dependency, the results suggest that the developmental timing of structural and microstructural features is shaped by complex, region-specific programs. Future region-focused or multivariate analyses may better elucidate the organizing principles that underlie the coupling, or uncoupling, of structural and microstructural maturation.

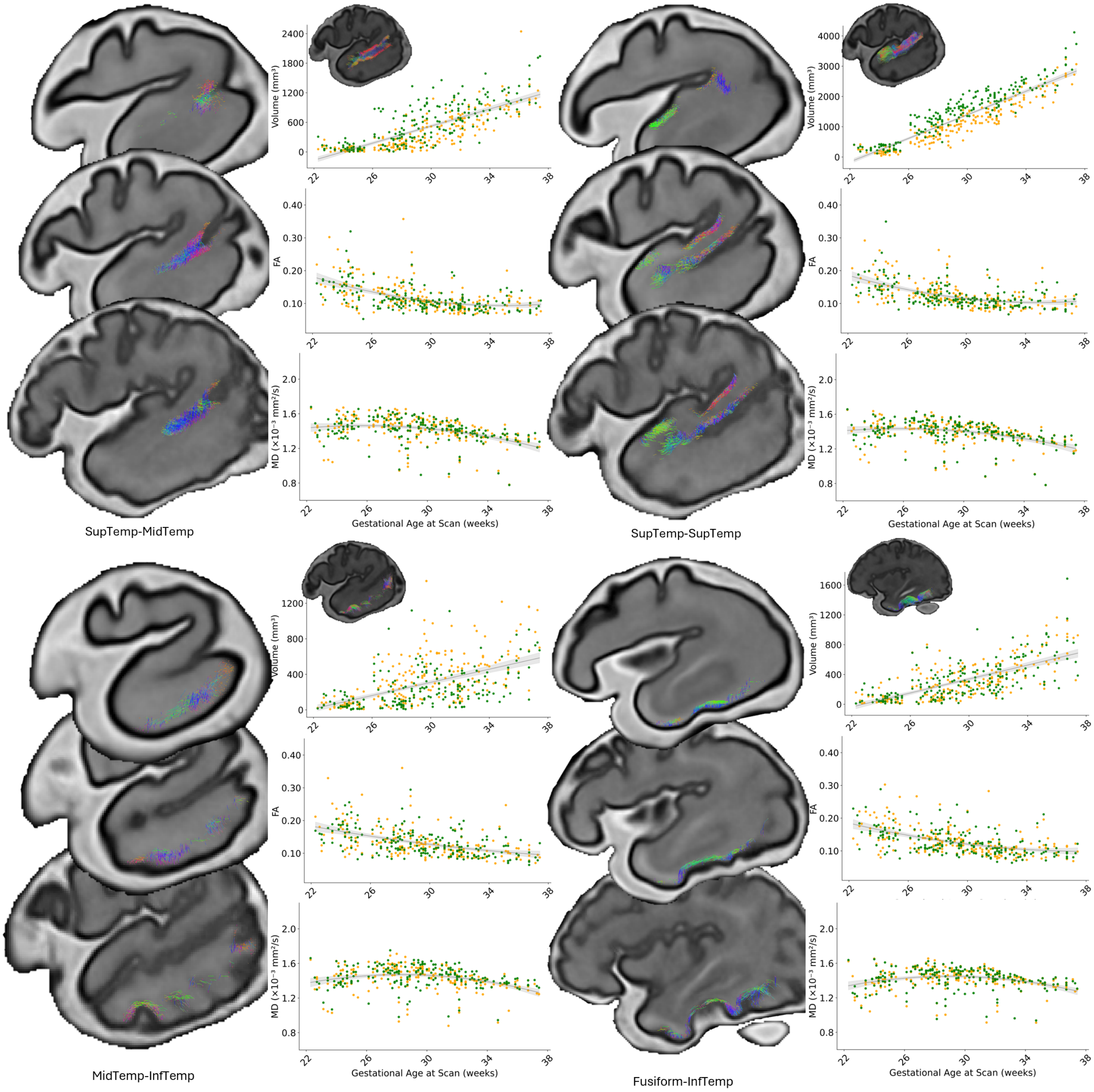


Figure A1: Four short association fiber (SAF) bundles in the temporal lobe are shown. SupTemp-MidTemp: Superior to Middle Temporal gyrus; SupTemp-SupTemp: SAFs within the Superior Temporal gyrus; MidTemp-InfTemp: Middle to Inferior Temporal gyrus; Fusiform-InfTemp: Fusiform to Inferior Temporal gyrus. For each bundle, developmental trajectories are visualized at 29, 32, and 35 GW, illustrating morphological changes over time. Bundle-specific measurements of volume (top; $mm^3$), free water-eliminated fractional anisotropy (middle; FA), and mean diffusivity (bottom; MD, $\times 10^{-3} mm^2/s$) are plotted across gestational age. FA and MD values represent the median of non-zero values within each bundle mask. Left (orange) and right (green) hemisphere measurements were jointly modeled. A linear model was used for volume, while DTI measures were fit using either linear or quadratic regression based on adjusted $R^2$. Shaded regions denote the 95% confidence intervals of the fitted trajectories, estimated via nonparametric bootstrapping (1000 iterations). Modeling conventions, regression methods, and color coding are consistent across all bundle figures (see Methods).

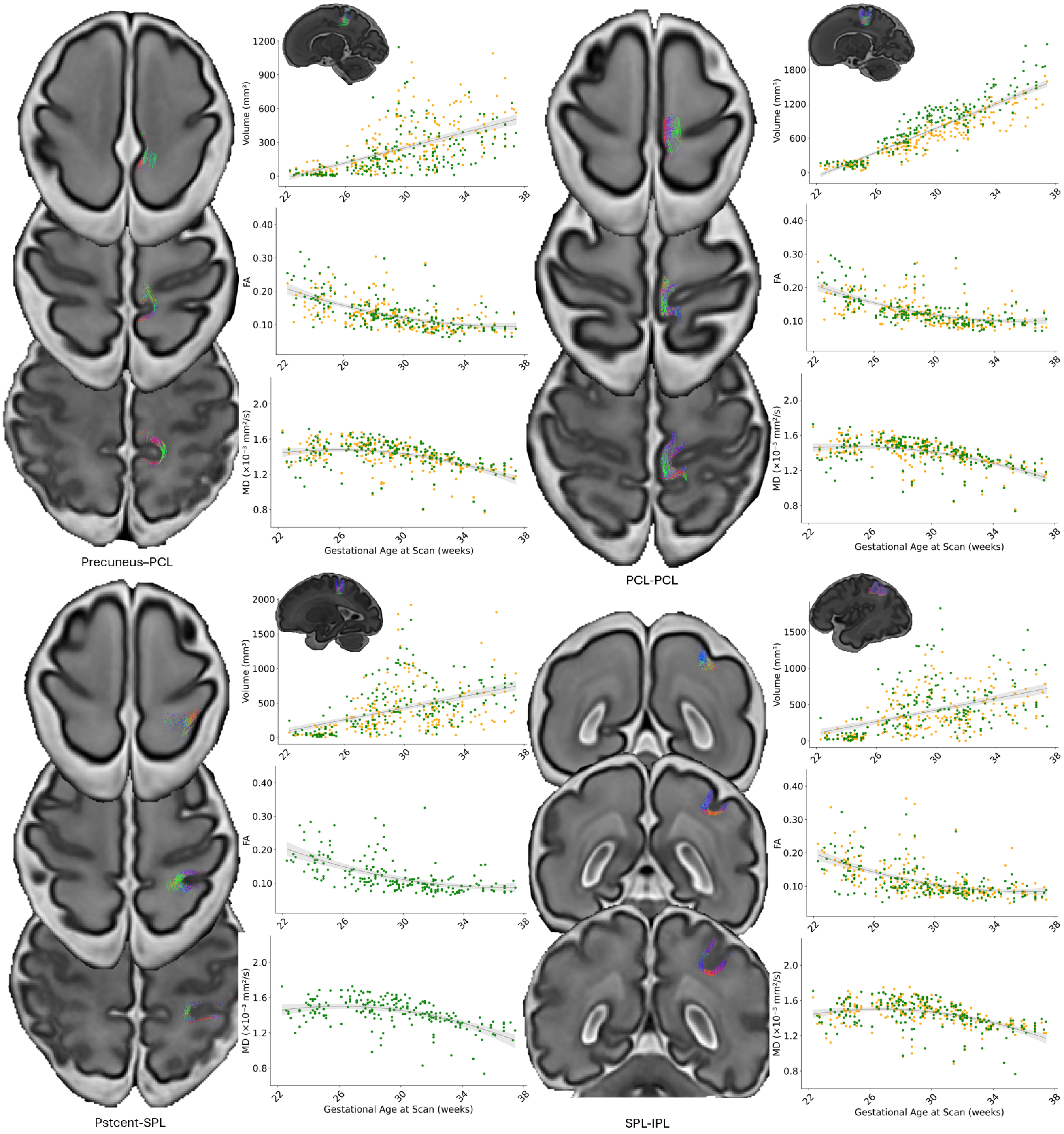


Figure A2: Four short association fiber (SAF) bundles in the the parietal lobe are shown. Precuneus-PCL: Precuneus to Paracentral Lobule; PCL-PCL: short association fibers within the Paracentral Lobule; Pstcent-SPL: Postcentral Gyrus to Superior Parietal Lobule; SPL-IPL: Superior to Inferior Parietal Lobule. For each bundle, developmental trajectories are visualized at 29, 32, and 35 gestational weeks, illustrating morphological changes over time. Bundle-specific measurements of volume (top; $mm^3$), free water-eliminated fractional anisotropy (middle; FA), and mean diffusivity (bottom; MD, $\times 10^{-3} mm^2/s$) are plotted across gestational age. FA and MD values represent the median of non-zero values within each bundle mask. Left (orange) and right (green) hemisphere measurements were jointly modeled. A linear model was used for volume, while DTI measures were fit using either linear or quadratic regression based on adjusted $R^2$. Shaded regions denote the 95% confidence intervals of the fitted trajectories, estimated via nonparametric bootstrapping (1000 iterations). Modeling conventions, regression methods, and color coding are consistent across all bundle figures (see Methods).

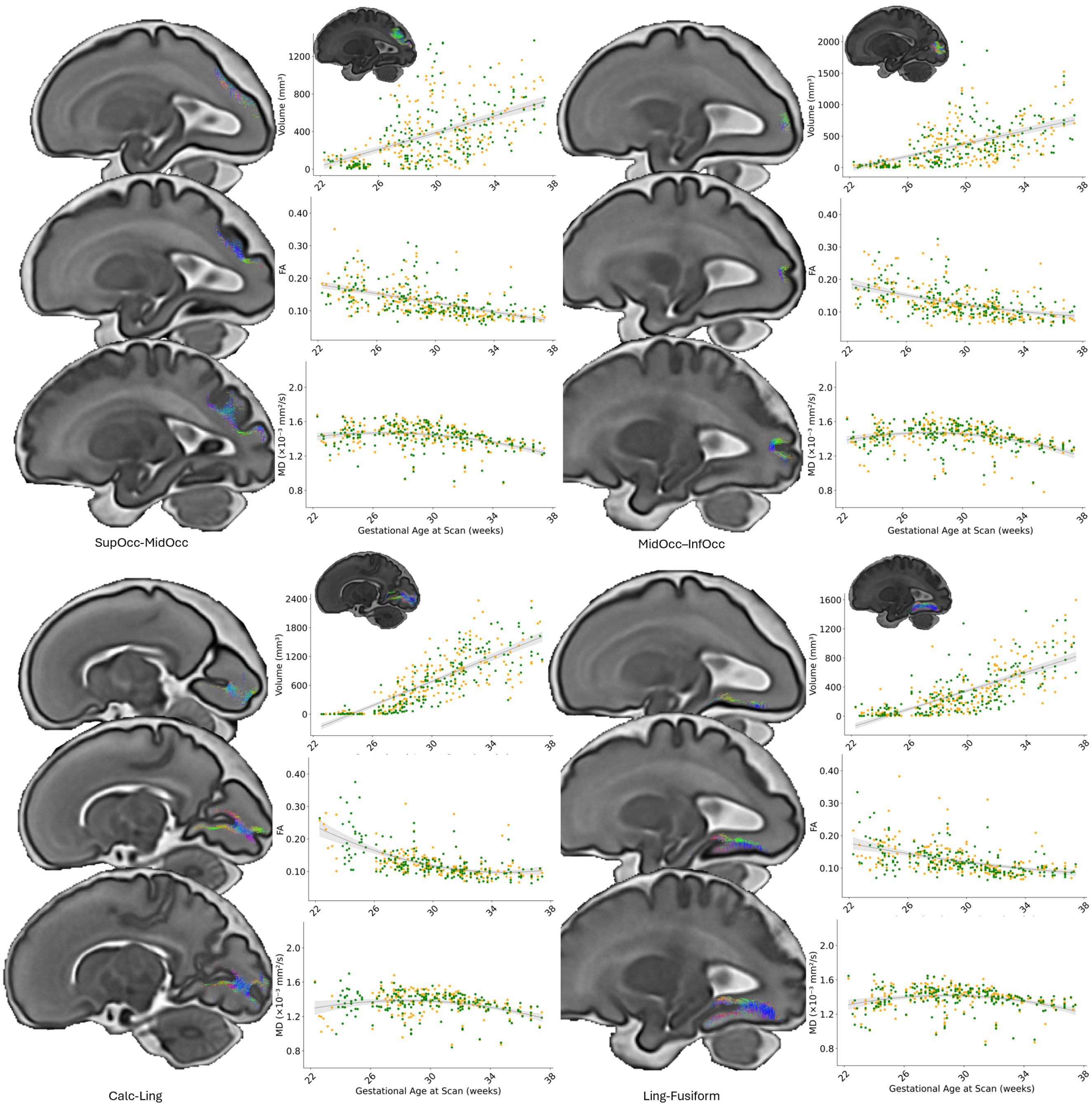


Figure A3: Four short association fiber (SAF) bundles in the the occipital lobe are shown. SupOcc-MidOcc: Superior to Middle Occipital Gyrus; MidOcc-InfOcc: Middle to Inferior Occipital Gyrus; Calc-Ling: Fibers connecting the Calcarine region (Cuneus side of the Calcarine sulcus) and the Lingual Gyrus; Ling-Fusiform: Lingual Gyrus to Fusiform Gyrus. For each bundle, developmental trajectories are visualized at 29, 32, and 35 gestational weeks, illustrating morphological changes over time. Bundle-specific measurements of volume (top; $mm^3$), free water-eliminated fractional anisotropy (middle; FA), and mean diffusivity (bottom; MD, $\times 10^{-3} mm^2/s$) are plotted across gestational age. FA and MD values represent the median of non-zero values within each bundle mask. Left (orange) and right (green) hemisphere measurements were jointly modeled. A linear model was used for volume, while DTI measures were fit using either linear or quadratic regression based on adjusted $R^2$. Shaded regions denote the 95% confidence intervals of the fitted trajectories, estimated via nonparametric bootstrapping (1000 iterations). Modeling conventions, regression methods, and color coding are consistent across all bundle figures (see Methods).

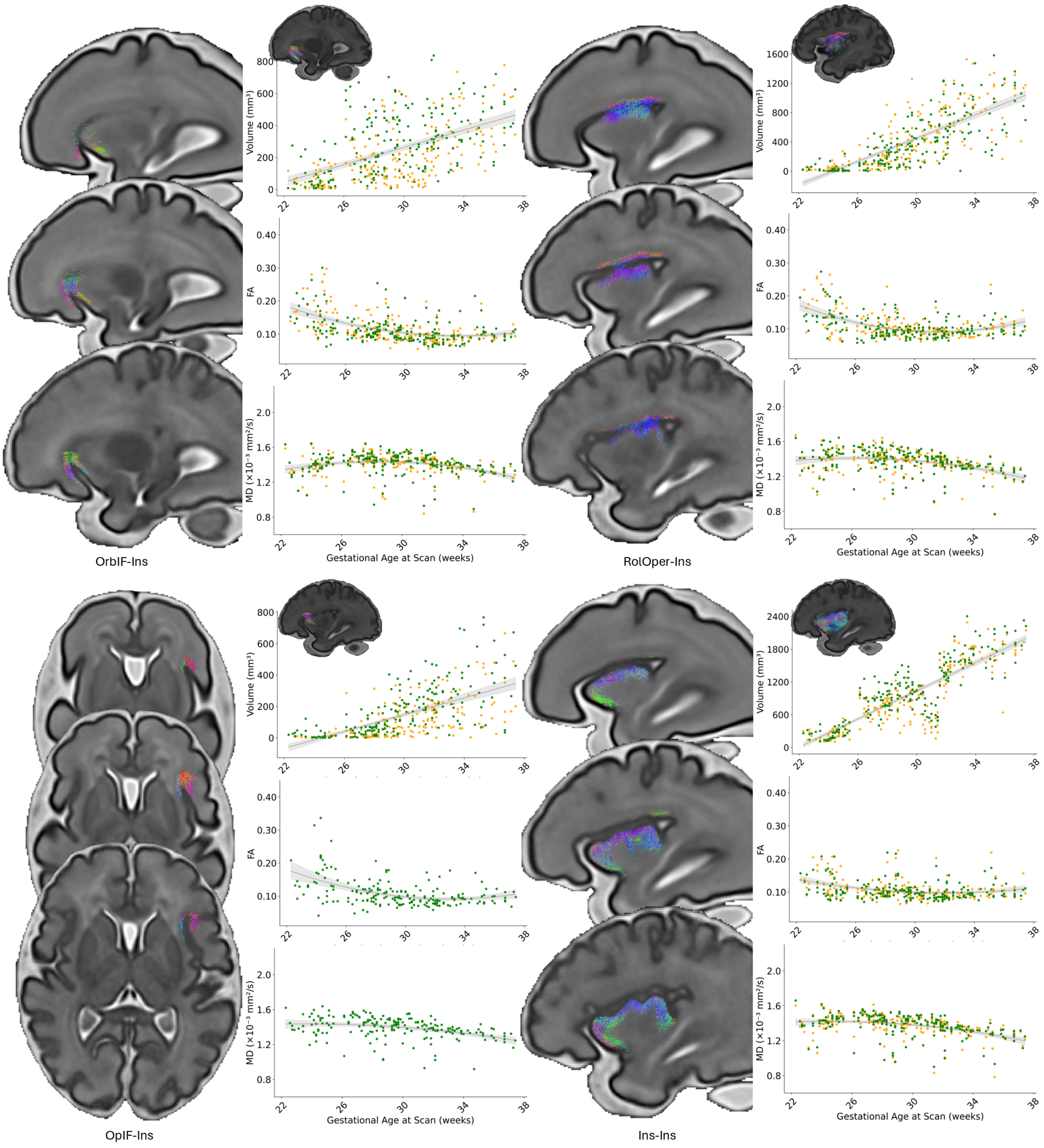


Figure A4: Four short association fiber (SAF) bundles in insular cortex are shown. OrbIF-Ins: Orbital Inferior Frontal Gyrus to Insular Cortex; RolOper-Ins: Rolandic Operculum to Insular Cortex; OpIF-Ins: Opercular Inferior Frontal Gyrus to Insular Cortex; Ins-Ins: short association fibers within the Insular Cortex. For each bundle, developmental trajectories are visualized at 29, 32, and 35 gestational weeks, illustrating morphological changes over time. Bundle-specific measurements of volume (top; $mm^3$), free water-eliminated fractional anisotropy (middle; FA), and mean diffusivity (bottom; MD, $\times 10^{-3} mm^2/s$) are plotted across gestational age. FA and MD values represent the median of non-zero values within each bundle mask. Left (orange) and right (green) hemisphere measurements were jointly modeled. A linear model was used for volume, while DTI measures were fit using either linear or quadratic regression based on adjusted $R^2$. Shaded regions denote the 95% confidence intervals of the fitted trajectories, estimated via nonparametric bootstrapping (1000 iterations). Modeling conventions, regression methods, and color coding are consistent across all bundle figures (see Methods).

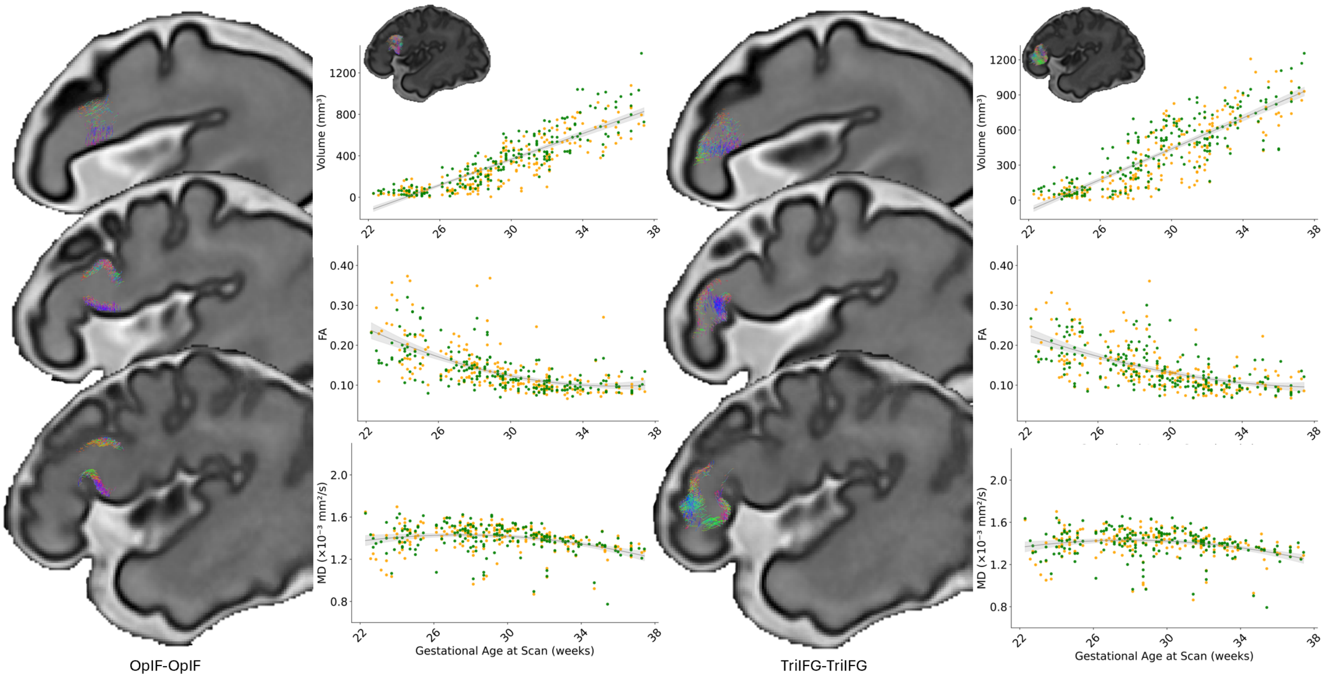


Figure A5: Additional short association fiber (SAF) bundles in the Frontal lobe. OpIF-OpIF: SAFs within the Opercular Inferior Frontal Gyrus; TriIFG-TriIFG: SAFs within the Triangular Inferior Frontal Gyrus. Bundle-specific measurements of volume (top; $mm^3$), free water-eliminated fractional anisotropy (middle; FA), and mean diffusivity (bottom; MD, $\times10^{-3}mm^2/s$) are plotted across gestational age. FA and MD values represent the median of non-zero values within each bundle mask. Left (orange) and right (green) hemisphere measurements were jointly modeled. A linear model was used for volume, while DTI measures were fit using either linear or quadratic regression based on adjusted $R^2$. Shaded regions denote the 95% confidence intervals of the fitted trajectories, estimated via nonparametric bootstrapping (1000 iterations). Modeling conventions, regression methods, and color coding are consistent across all bundle figures (see Methods).

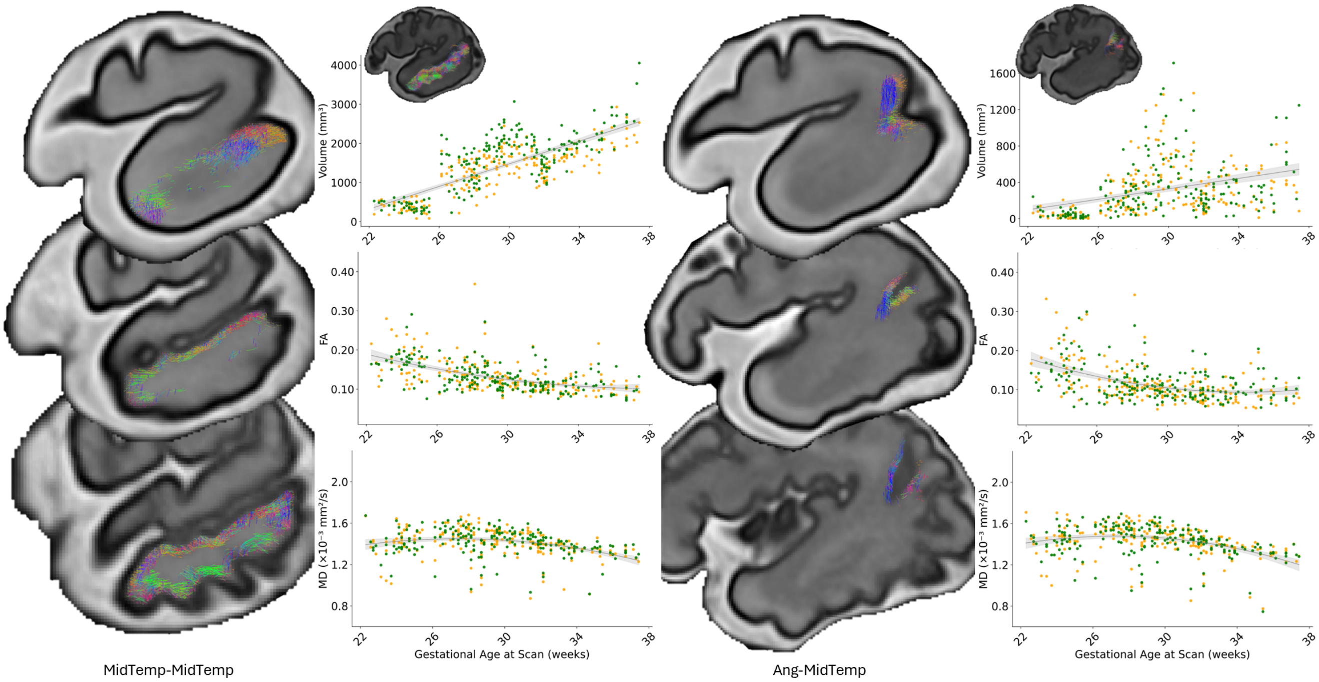


Figure A6: Additional short association fiber (SAF) bundles in the Temporal lobe. MidTemp-MidTemp: SAFs within the Middle Temporal Gyrus; Ang-MidTemp: Angular Gyrus to Middle Temporal Gyrus. Bundle-specific measurements of volume (top; $mm^3$), free water-eliminated fractional anisotropy (middle; FA), and mean diffusivity (bottom; MD, $\times 10^{-3} mm^2/s$) are plotted across gestational age. FA and MD values represent the median of non-zero values within each bundle mask. Left (orange) and right (green) hemisphere measurements were jointly modeled. A linear model was used for volume, while DTI measures were fit using either linear or quadratic regression based on adjusted $R^2$. Shaded regions denote the 95% confidence intervals of the fitted trajectories, estimated via nonparametric bootstrapping (1000 iterations). Modeling conventions, regression methods, and color coding are consistent across all bundle figures (see Methods).

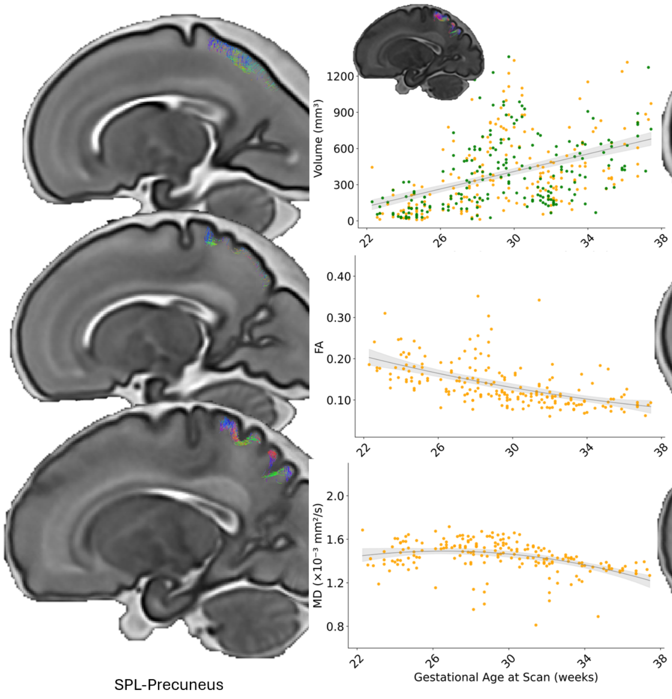


Figure A7: Additional short association fiber (SAF) bundles in the Parietal lobe. SPL-Precuneus: Superior Parietal Lobule to Precuneus; Precuneus-Precuneus: SAFs within the Precuneus. Bundle-specific measurements of volume (top; $mm^3$), free water-eliminated fractional anisotropy (middle; FA), and mean diffusivity (bottom; MD, $\times 10^{-3} mm^2/s$) are plotted across gestational age. FA and MD values represent the median of non-zero values within each bundle mask. Left (orange) and right (green) hemisphere measurements were jointly modeled. A linear model was used for volume, while DTI measures were fit using either linear or quadratic regression based on adjusted $R^2$. Shaded regions denote the 95% confidence intervals of the fitted trajectories, estimated via nonparametric bootstrapping (1000 iterations). Modeling conventions, regression methods, and color coding are consistent across all bundle figures (see Methods).

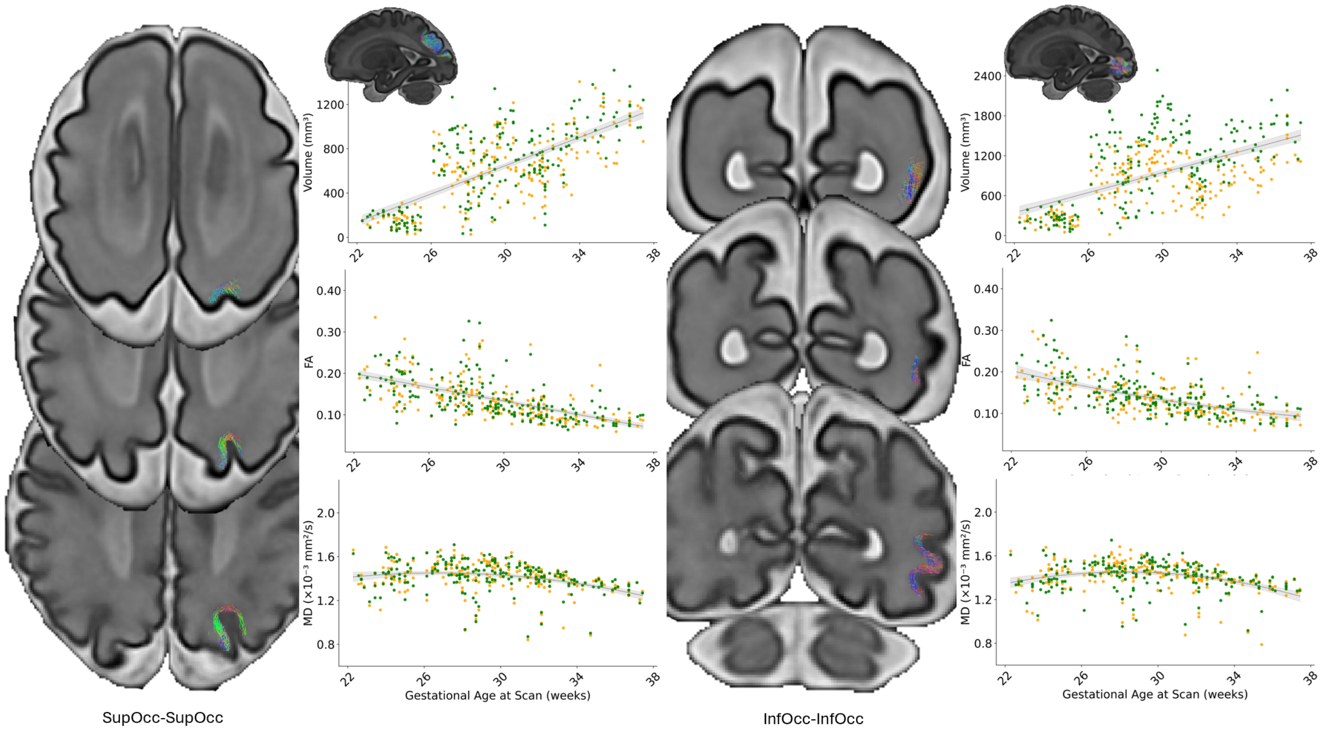


Figure A8: Additional short association fiber (SAF) bundles in the Occipital lobe. SupOcc-SupOcc: SAFs within the Superior Occipital Gyrus; InfOcc-InfOcc: SAFs within the Inferior Occipital Gyrus. Bundle-specific measurements of volume (top; $mm^3$), free water-eliminated fractional anisotropy (middle; FA), and mean diffusivity (bottom; MD, $\times 10^{-3} mm^2/s$) are plotted across gestational age. FA and MD values represent the median of non-zero values within each bundle mask. Left (orange) and right (green) hemisphere measurements were jointly modeled. A linear model was used for volume, while DTI measures were fit using either linear or quadratic regression based on adjusted $R^2$. Shaded regions denote the 95% confidence intervals of the fitted trajectories, estimated via nonparametric bootstrapping (1000 iterations). Modeling conventions, regression methods, and color coding are consistent across all bundle figures (see Methods).

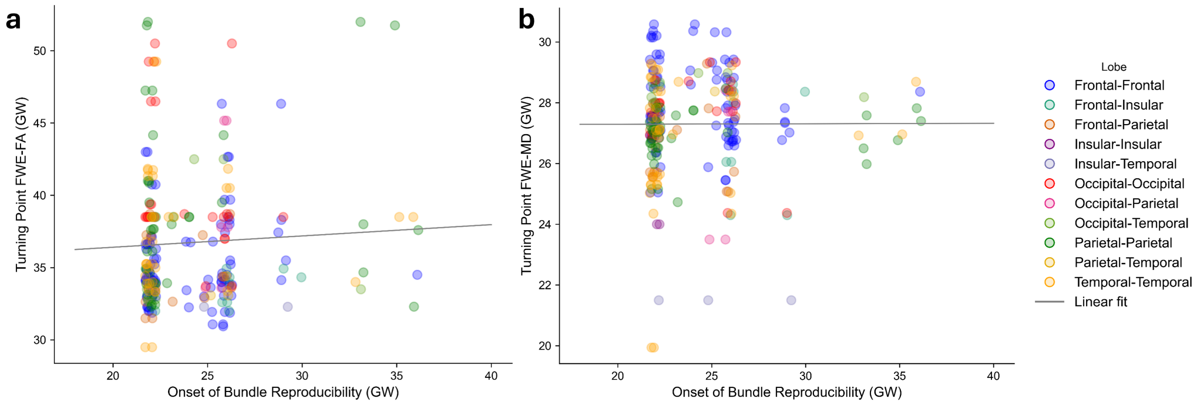


Figure A9: Relationship between the anatomical onset and microstructural turning points of short association fiber bundles. Scatter plots show the age of anatomical reproducibility (x-axis) versus the turning points of free-water–eliminated fractional anisotropy (FA, **a**) and mean diffusivity (MD, **b**) (y-axis) across all bundles. Each point represents one bundle, colored by lobar classification. Fitted linear regression lines are shown for reference (FA: $y = 34.84 + 0.078x$, p-value=0.418, $R^2$=0.0028; MD: $y = 27.26 + 0.0015x$, p-value=0.968, $R^2$ <0.0001). These results indicate no significant linear relationship between anatomical onset and the FA or MD turning points. The observed distribution instead suggests regional clustering without a strong monotonic trend.

### Methods

**Algorithm A1** Hierarchical extraction and refinement of short association fiber bundles

**Input:** Preprocessed dMRI data, tissue segmentation, parcellation atlas
Compute FOD, DTI, sharpened dODF, FA, and MD maps
Constrain tractography to short-range fibers within the superficial white matter system
Perform gyri-based bundle extraction and group-level refinement:

- Extract all streamlines with both endpoints in a pair of gyri using cortical parcellation
- For each bundle:
  - Perform streamline clustering and compute confidence index per streamline
  - Prune streamlines with confidence index below the 3rd or 5th percentile
  - Exclude bundles with fewer than 100 streamlines
- For each age group:
  - Spatially register individual data to the age-specific dHCP fetal atlas
  - Compute the Dice coefficient between individual and group-average bundle masks
  - Exclude bundles with fewer than $N$ valid Dice scores (Dice $> 0.1$, $N >$ half the group size)
  - Iteratively remove subjects whose exclusion most improves the average Dice score
    - Stop when $\Delta\widehat{\text{Dice}} < 0.005$, $p > 0.01$, or remaining group size $< 5$
- Retain bundles that pass quality control across all age groups
- Update group-average bundle masks using retained subjects

**Output:** Refined, high-confidence superficial bundles across gestational ages

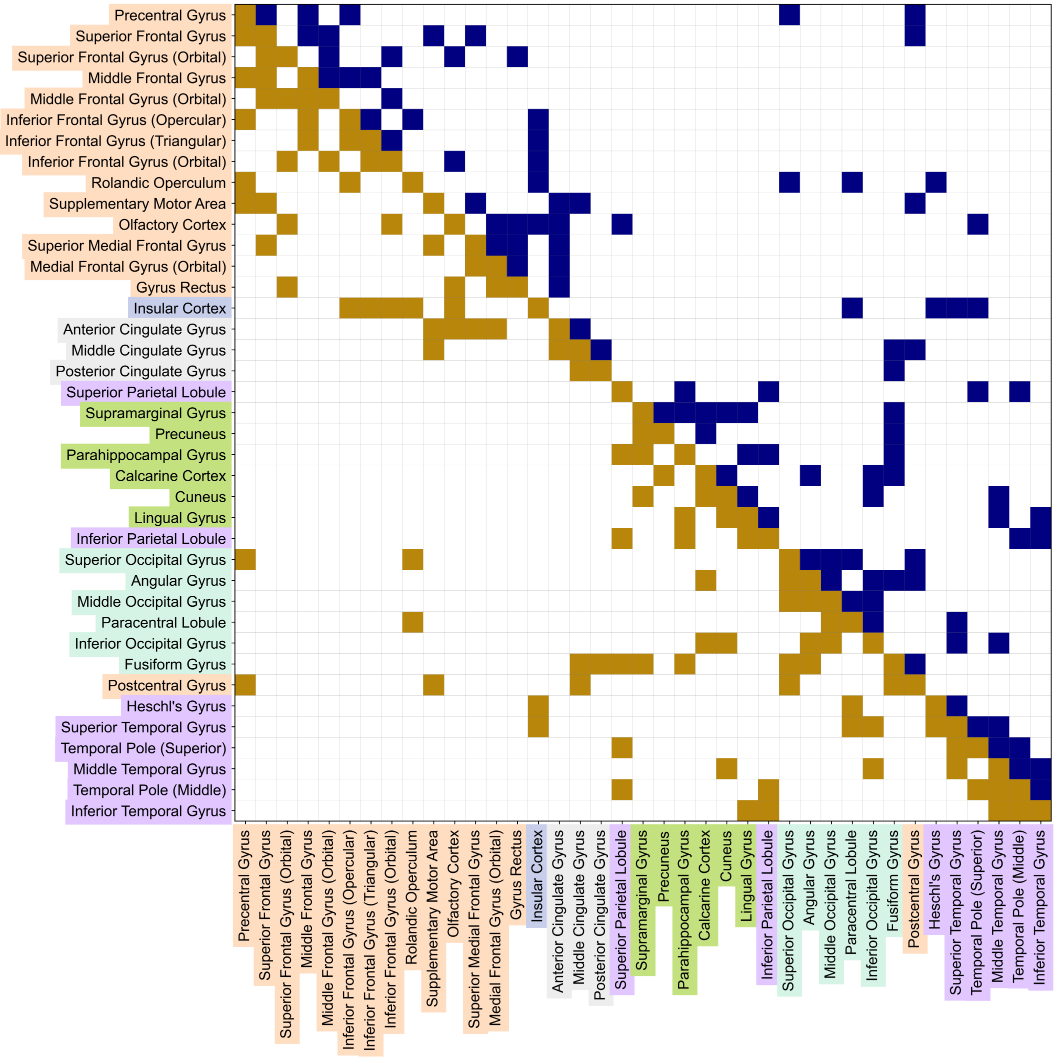


Figure A10: Identified gyri-level short-range structural connections at 36 gestational weeks. The lower triangle (golden brown) represents connections identified in the left hemisphere, while the upper triangle (excluding the diagonal; blue) represents those in the right hemisphere. The robust presence of connections is not necessarily symmetric between hemispheres. Diagonal elements denote intra-gyri connections, which are present in both hemispheres, Axis labels correspond to cortical regions, color-coded by their anatomical locations (lobes).

Table A1: Gyri names, anatomical regions, and emergence stage according to their first recognizable age (Chi et al., 1977). We refer "Early" to those identified <26 weeks (wk) of gestation, "Mid" between 26-31 weeks, and "Late" ≥ 32 weeks. Left and right hemispheres are not distinguished in the gyrus names. Asterisks mark gyri without a direct definition in Chi et al. (1977); their folding stage is estimated based on the timing of the most closely corresponding classical gyrus. For example, the Superior Medial Frontal Gyrus is considered part of the callosomarginal gyrus, which becomes defined at approximately 29 wk.

| **Gyrus Name** | **Abbreviation** | **Lobe** | **Emergence Stage** |
|---|---|---|---|
| Precentral Gyrus | PreC | Frontal Lobe | Early (≈ 24 wk) |
| Superior Frontal Gyrus | SupF | Frontal Lobe | Early (≈ 25 wk) |
| Superior Frontal Gyrus (Orbital) | OrbSF | Frontal Lobe | Late (≈ 36 wk) |
| Middle Frontal Gyrus | MidF | Frontal Lobe | Mid (≈ 27 wk) |
| Middle Frontal Gyrus (Orbital) | OrbMidF | Frontal Lobe | Late (≈ 36 wk) |
| Inferior Frontal Gyrus (Opercular) | OpIF | Frontal Lobe | Mid (≈ 28 wk) |
| Inferior Frontal Gyrus (Triangular) | TriIFG | Frontal Lobe | Mid (≈ 28 wk) |
| Inferior Frontal Gyrus (Orbital) | OrbIF | Frontal Lobe | Mid (≈ 28 wk) |
| Rolandic Operculum | RolOper | Frontal Lobe | Mid* (≈ 28-29 wk) |
| Supplementary Motor Area | SMA | Frontal Lobe | Mid* (≈ 29 wk) |
| Olfactory Cortex | Olf | Frontal Lobe | Early (≈ 16 wk) |
| Superior Medial Frontal Gyrus | MedSupF | Frontal Lobe | Mid* (≈ 29 wk) |
| Medial Frontal Gyrus (Orbital) | OrbMF | Frontal Lobe | Mid (≈ 28 wk) |
| Gyrus Rectus | Rect | Frontal Lobe | Early (≈ 16 wk) |
| Insular Cortex | Ins | Insular Cortex | Early (≈ 18 wk) |
| Anterior Cingulate Gyrus | AntCng | Frontal Lobe | Early (≈ 18 wk) |
| Middle Cingulate Gyrus | MidCng | Parietal Lobe | Early (≈ 18 wk) |
| Posterior Cingulate Gyrus | PostCng | Parietal Lobe | Early (≈ 18 wk) |
| Calcarine Cortex | Calc | Occipital Lobe | Early (≈ 16 wk) |
| Cuneus | Cuneus | Occipital Lobe | Mid (≈ 27 wk) |
| Lingual Gyrus | Ling | Occipital Lobe | Mid (≈ 27 wk) |
| Superior Occipital Gyrus | SupOcc | Occipital Lobe | Mid (≈ 27 wk) |
| Middle Occipital Gyrus | MidOcc | Occipital Lobe | Mid (≈ 27 wk) |
| Inferior Occipital Gyrus | InfOcc | Occipital Lobe | Mid (≈ 27 wk) |
| Postcentral Gyrus | Pstcent | Parietal Lobe | Early (≈ 25 wk) |
| Superior Parietal Lobule | SPL | Parietal Lobe | Mid (≈ 26 wk) |
| Inferior Parietal Lobule | IPL | Parietal Lobe | Mid (≈ 26 wk) |
| Supramarginal Gyrus | SMG | Parietal Lobe | Mid (≈ 28 wk) |
| Angular Gyrus | Ang | Parietal Lobe | Mid (≈ 28 wk) |
| Precuneus | Precuneus | Parietal Lobe | Mid* (≈ 28-32 wk) |
| Paracentral Lobule | PCL | Frontal Lobe | Late (≈ 34-35 wk) |
| Parahippocampal Gyrus | ParaHip | Temporal Lobe | Early (≈ 23 wk) |
| Fusiform Gyrus | Fusiform | Temporal Lobe | Mid (≈ 27 wk) |
| Heschl's Gyrus | TransTMP | Temporal Lobe | Mid (≈ 31 wk) |
| Superior Temporal Gyrus | SupTemp | Temporal Lobe | Early (≈ 23 wk) |
| Temporal Pole (Superior) | SupTP | Temporal Lobe | Late* |
| Middle Temporal Gyrus | MidTemp | Temporal Lobe | Mid (≈ 26 wk) |
| Temporal Pole (Middle) | MidTP | Temporal Lobe | Late* |
| Inferior Temporal Gyrus | InfTemp | Temporal Lobe | Mid (≈ 30 wk) |
